# A New ‘Comprehensive’ Annotation of Leucine-Rich Repeat-Containing Receptors in Rice

**DOI:** 10.1101/2021.01.29.428842

**Authors:** Céline Gottin, Anne Dievart, Marilyne Summo, Gaëtan Droc, Christophe Périn, Vincent Ranwez, Nathalie Chantret

**Affiliations:** AGAP, Univ Montpellier, CIRAD, INRAE, Institut Agro, Montpellier, France; CIRAD, UMR AGAP, F-34398 Montpellier, France

## Abstract

Rice plays an essential food security role for more than half of the world’s population. Obtaining crops with high levels of disease resistance is a major challenge for breeders, especially today given the urgent need for agriculture to be more sustainable. Plant resistance genes are mainly encoded by three large Leucine-Rich Repeat (LRR)-containing receptor (LRR-CR) subfamilies: the LRR-Receptor-Like Kinase (RLK), LRR-Receptor-Like Protein (RLP) and Nucleotide-binding LRR Receptor (NLR) subfamilies. Using LRRprofiler, a pipeline we developed to annotate and classify those proteins, we compared three publicly available annotations of the rice Nipponbare reference genome. The extended discrepancies we observed for LRR-CR gene models led us to perform in-depth manual curation of their annotations while paying special attention to nonsense mutations. We then transferred this manually curated annotation to Kitaake, a Nipponbare closely related cultivar, using an optimised strategy. Here we discuss the breakthrough achieved by manual curation when comparing genomes and, in addition to ‘functional’ and ‘structural’ annotations, we propose the community to adopt this new approach, which we call ‘comprehensive’ annotation. The resulting data are crucial for further studies on the natural variability and evolution of LRR-CR in order to promote their use in breeding future resilient varieties.

## Introduction

Modern agriculture is at a critical juncture as the world’s population continues to grow while there is a call for a shift away from chemical input applications to deal with current environmental issues. Crop pest and pathogen susceptibility is one of the main causes of annual crop yield loss (FAO, 2018; Savary et al., 2019) and declining produce quality. Despite awareness on the harmful environmental impacts, massive pesticide use is still a common means to prevent plant diseases today. Studying and understanding plant disease resistance and the underlying evolutionary mechanisms is of utmost importance to make effective widespread use of known sources of resistance through specific breeding programs, while also promoting new resistance engineering for crop sustainability (Bailey-Serres et al., 2019; Tamborski and Krasileva, 2020). The elucidation of resistance mechanisms in plants has highlighted a trove of resistance genes to combat the high and evolving genetic diversity of plant pathogens. The Leucine-Rich Repeat (LRR)-containing receptor (LRR-CR) is at the forefront among these genes. LRR-CRs share the common structural and functional LRR domain. This domain is composed of two to more than 30 repetitions of a ∼24 amino acid motif characterised by a conserved skeleton composed mostly of leucine residues (Kajava, 1998; Bella et al., 2008; Kajava, 2012; Matsushima and Miyashita, 2012). LRR-CRs are classified in three main gene subfamilies: LRR Receptor-Like Kinase (LRR-RLK), LRR Receptor-Like Protein (LRR-RLP) and Nucleotide-Binding site LRR (NBS-LRR or NLR) (Sekhwal et al., 2015; Han, 2019) (**Figure 1**). LRR-RLKs and LRR-RLPs are referred to as pattern-recognition receptors (PRRs). These transmembrane receptors have an extracellular LRR domain and an intracellular domain. The latter is a kinase domain for LRR-RLK and a short cytoplasmic tail for LRR-RLP. Note that LRR-RLKs are known intercellular communication players, thus they are also involved in other biological processes such as development. NLRs are intracellular receptors composed of an NB-ARC domain followed by the LRR domain (Sekhwal et al., 2015; Burdett et al., 2019; Tamborski and Krasileva, 2020; Xiong et al., 2020).

**Figure 1.**
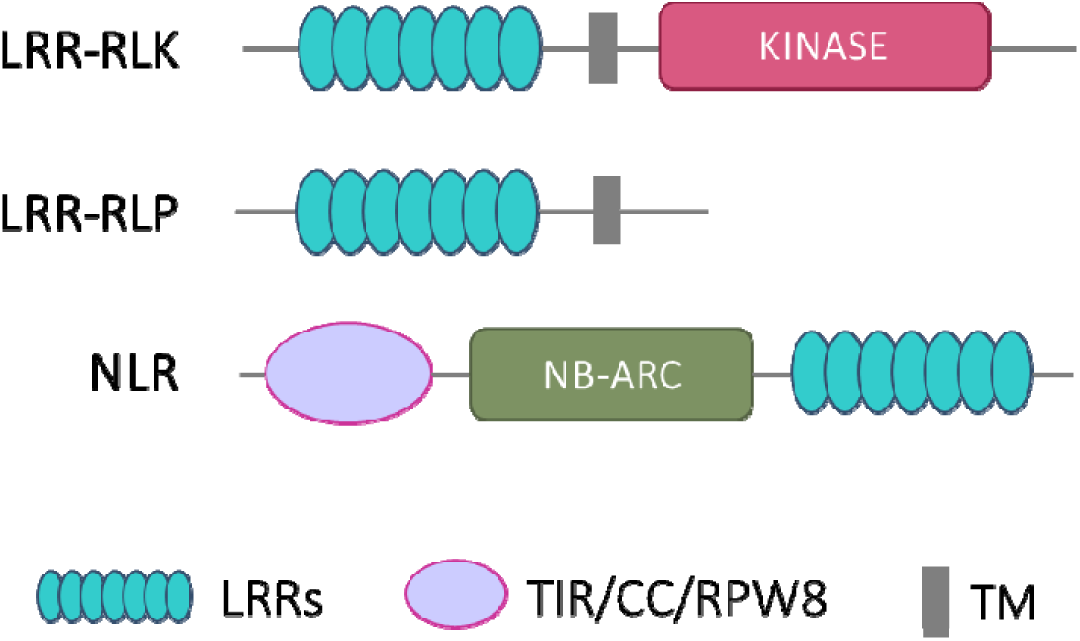
Schematic protein structure of the three LRR-CR subfamilies: LRR-RLK, LRR-RLP and NLR. TM, transmembrane domain; CC, coiled coil domain; TIR, toll-interleukin receptor; RPW8, resistance to powdery mildew 8 domain.

Over the past few decades, advances in sequencing have provided the research community with an ever-increasing number of complete genomes. These resources have made it possible to revisit gene evolution at the level of entire families and on different evolutionary time scales. LRR-CR genes have been inventoried in many angiosperm genomes, and their numbers have also been compared in a phylogenetic framework to shed light on their evolutionary dynamics (Wang et al., 2011; Sharma and Pandey, 2015; Fischer et al., 2016; Dufayard et al., 2017). A large proportion of LRR-CR genes are thought to evolve through a so called birth-and-death model (Michelmore and Meyers, 1998; Richter and Ronald, 2000; Nei and Rooney, 2005; McDowell and Simon, 2006). In this model, the gene copy number expands by recurrent duplication events and duplicated copies can then follow different evolutionary pathways such as keeping the original function, acquiring a new one (neo-functionalization) or, more frequently, undergoing a non-functionalization process by accumulating nonsense mutations (Leister, 2004; Innan and Kondrashov, 2010). This model explains why LRR-CR genes are found in multiple copies, often organized in large gene clusters, with some genes no longer being functional (Meyers et al., 2003; Mizuno et al., 2020).

Comparative genomic studies have led to considerable progress in understanding the evolutionary dynamics of LRR-CR gene families, but these studies are highly dependent on the accuracy of annotation procedures. Given the increasing avalanche of sequence data, the most commonsense approach is to rely on automatic annotation. Gene and protein sequence annotation are thus crucial and the focus of considerable effort. Structural gene annotation is geared towards identifying coding sequences within genomic data and documenting the associated gene features (e.g. introns, exons, UTRs) (Wilming and Harrow, 2009). The most widely used structural annotation pipelines (e.g. Ensembl pipeline for gene annotation (Aken et al., 2016), Augustus (Stanke and Waack, 2003) and Gnomon (https://www.ncbi.nlm.nih.gov/genome/annotation_euk/gnomon/)) rely (i) on *ab initio* gene structure determination according to rules learned on pre-existing annotations, and/or (ii) on comparative approaches i.e. using sequence homology with available RNAseq data and/or with a closely related annotated genome. Those methods allow large-scale studies with standardized approaches, yet they are not completely reliable, especially for complex multigene families. Indeed, repetitions are known to impair gene annotations (Fawal et al., 2014; Bayer et al., 2018) and there are also genome construction issues (Torresen et al., 2019). The difficulty is twofold in the case of LRR-CRs: several similar genes are present in the genome due to gene duplication events, while each gene contains several similar domains due to the repetitive structure of the LRR domain. Automatic annotation and classification of LRR-CRs is thus especially challenging. For example, although multiple studies have reported that there are more than 800 LRR-CR loci in the Nipponbare rice variety, the number of genes per subfamily is variable, e.g. 374 to 498 NLR proteins (Zhou et al., 2004; Li et al., 2010; Li et al., 2016; Shao et al., 2016; Stein et al., 2018), 292 to 332 LRR-RLKs (Hwang et al., 2011; Sun and Wang, 2011; Dufayard et al., 2017) and 90 LRR-RLPs (Fritz-Laylin et al., 2005). These variations are to a large extent linked to the annotation version chosen for the analysis and to the decision rules for gene detection and classification. Scientists sometimes perform manual curation of gene annotations to limit these uncertainties and achieve high quality comprehensive analyses, as for Arabidopsis and tomato NLR genes (Meyers et al., 2003; Jupe et al., 2013; Van de Weyer et al., 2019) or Nipponbare rice LRR-RLK genes (Sun and Wang, 2011).

Rice was the first monocotyledon plant to have its genome entirely sequenced and three different annotations of its reference genome, *O. sativa* ssp. *japonica* cv. Nipponbare (Kawahara et al., 2013), are currently available: one from the Michigan State University Rice Genome Annotation Project (hereafter named MSU) (Yuan et al., 2003), one from the National Center for Biotechnology Information (NCBI) and the current reference one from the Rice Annotation Project of the International Rice Genome Sequencing Project (IRGSP) (Sakai et al., 2013). We first implemented the LRRprofiler pipeline to compare them with regard to the LRR-CR repertoire. This program builds subfamily- and genome-specific LRR Hidden Markov Model (HMM) profiles, detects LRR-CR genes that contain LRR motifs, and accurately locates all LRR motifs within these genes. We ran the LRRprofiler pipeline in parallel on the three rice annotations and found that they greatly differed in terms of the number of LRR-CR genes and their structural annotations. We therefore performed a manual curation of the whole Nipponbare LRR-CR repertoire. We decided not to force gene models to have to contain all expected features in functional genes but rather, when appropriate, to supplement the gene models with sequence fragments undoubtedly derived from LRR-CR encoding genes. In turn, we provided objective information, i.e. whether the gene models were canonical or non-canonical. To be qualified as canonical, a gene model had to fulfil all of the following conditions: presence of a start codon, presence of a terminal stop codon, absence of an in-frame stop codon, absence of frameshifts, and absence of unexpected intron splicing sites. Conversely, any gene violating at least one of these constraints was qualified as non-canonical. Finally, we also propose a strategy to transfer these manually curated LRR-CR gene annotations to Kitaake, the closest related japonica genome whose sequence is available (Jain et al., 2019). We then studied observed variations in gene numbers and LRR motifs between Nipponbare and Kitaake genotypes when using available automatic annotations and our manually curated annotations (hereafter referred to as ‘comprehensive’). This comparison demonstrated how erroneous conclusions can readily be drawn when relying solely on automatic structural and functional annotations for this complex gene family. The curated comprehensive LRR-CR annotation introduced in this article is available online through a dedicated website (https://rice-genome-hub.southgreen.fr/content/geloc).

## Results

### Inconsistencies among three publicly available Nipponbare rice LRR-CR annotations

We used LRRprofiler, a newly developed pipeline (see Material and Methods and **Supplemental Methods**), to identify, annotate and classify into gene subfamilies the LRR-CR protein sequences of the three publicly available Nipponbare proteomes (MSU, IRGSP, NCBI). The total number of LRR-CRs identified varied markedly according to the annotation: we identified 1,186 LRR-CRs in the MSU predicted proteome, 981 in that of IRGSP and 1,025 in that of NCBI (**Table 1**). The distribution patterns of these proteins in the different subfamilies also varied according to the annotations. For instance, the number of predicted genes fluctuated less for the RLP subfamily than for the NLR subfamily for which 64% more NLRs were detected in the MSU proteome (405 proteins) compared to the IRGSP proteome (260 proteins).

**Table 1.** Number of LRR-CR sequences in the predicted proteomes from three publicly available annotations for the Nipponbare rice reference genome. Sequences were identified and classified into subfamilies using the LRRprofiler pipeline.

|  | Total | LRR-RLKs | LRR-RLPs | NLRs | Others* |
| --- | --- | --- | --- | --- | --- |
| IRGSP | 981 | 237 (24.2%) | 160 (16.3%) | 260 (26.5%) | 324 (33.0%) |
| MSU | 1,186 | 329 (27.7%) | 140 (11.8%) | 405 (34.2%) | 312 (26.3%) |
| NCBI | 1,025 | 305 (29.8%) | 122 (11.9%) | 339 (33.1%) | 259 (25.2%) |

\* Others refer here to F-box-LRR and unclassified (UC) sequences

We assessed whether the identified genes were at the same genomic location or not by measuring the overlap of the three predicted gene sets. We conducted a similar analysis on nine transcription factor (TF) subfamilies, for which we assumed that the annotation process would be easier since they had a more conserved structure and, although having undergone expansion events, were not evolving under a birth-and-death model (Lai et al., 2020). The TF dataset contained between 874 and 1,041 genes, according to the annotations, and this number was similar to that of LRR-CR. The percentage of loci for which a gene model was present in all three annotations was 52.9% for LRR-CR genes and 69.5% for TF (**Figure 2A**), indicating that the three annotations were more congruent for TF genes. Moreover, the percentage of loci in which only one annotation detected a gene was 19.5% for LRR-CR genes, compared to only 12% for TF genes.

**Figure 2.**
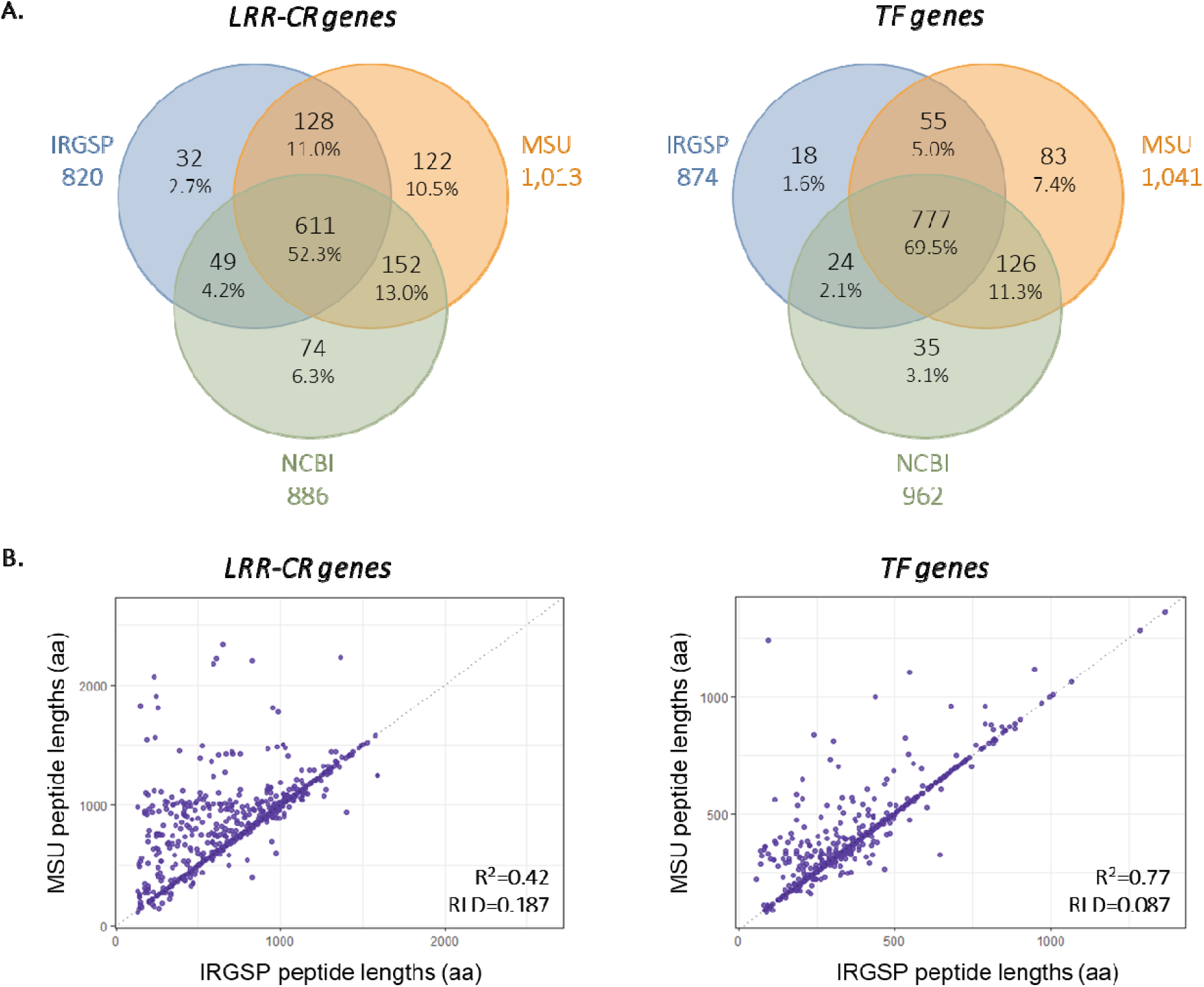
Comparison of MSU, IRGSP and NCBI publicly available annotations for the Nipponbare rice reference genome for two types of genes: LRR-containing receptors (LRR-CR) and transcription factors (TF). **(A)** Venn diagrams representing the number of overlapping gene models for LRR-CR and TF between the MSU, IRGSP and NCBI annotations. To be considered as overlapping, gene models from two (or three) different annotations should have at least one nucleotide in common (overlapping loci). **(B)** Dot plots representing the polypeptide length in amino acids (aa) for genes predicted by both IRGSP and MSU annotations. On the left, LRR-CRs and on the right, TFs. The R^2^ and the average relative length difference (RLD) are given at the bottom right for each gene family. See **Supplemental Figure 1** for all pairwise comparisons between IRGSP, MSU and NCBI.

Even when a gene was predicted by the different annotations, the predicted structure of the gene sometimes varied between predictions. One way to address this issue is to compare the length of predicted proteins for genes positioned at the same locus. Note that this is a conservative approach. Indeed, although a predicted protein length difference between two gene models indicated that the gene models differed, the reverse was not true as identical predicted protein lengths did not guarantee that the gene models were identical. A comparison of predicted protein lengths for all LRR-CR gene pairs located at the same locus but predicted by two different annotations is presented in **Figure 2B** and **Supplemental Figure 1**. Here again, the number of genes with a difference in the predicted protein length highlighted a substantial annotation discrepancy. This difference was greater for LRR-CR genes than for TF genes. As an example, when IRGSP and MSU were compared, the average difference between the predicted product sizes was 18.7% for LRR-CR loci (with a R^2^ of only 0.42) whereas this difference was only 8.6% for TF loci (with a much higher R^2^ of 0.77) (**Figure 2B**). These results highlight the extent to which annotations generally differ, but more particularly for LRR-CR gene subfamilies. These comparisons also showed that LRR-CR genes predicted by IRGSP were generally shorter than those predicted by MSU or NCBI at the same locus (**Figure 2B** and **Supplemental Figure 1**).

### Manual re-annotation of LRR-CR encoding loci in the Nipponbare rice genome

Here we provide a brief description of the procedure we followed to manually curate LRR-CR annotations (**Figure 3A**). Note above all that, for traceability sake, the procedure retained one of the three proposed gene models as much as possible. For a given locus, we first selected one of the gene models among the available annotations based on the completeness of the predicted protein. We then applied our expertise to the selected gene model by combining protein and nucleic data. At the protein level, we checked that all of the expected domains for each subfamily were present (e.g. LRRs and TM for LRR-RLPs and LRR-RLKs, or NB-ARC for NLRs), in the right order, with the expected length and inter-domain intervals. Protein domain information was particularly useful for detecting potential gene fusion and fission. At the nucleic level, we examined: (i) if the gene models had the expected structure (intron/exon), (ii) if nearby open reading frames (ORF) belonging to LRR-CR encoding sequences were present, and (iii) if the gene models included suspicious introns, such as short introns thereby enabling the gene to sidestep stop codons or frameshifts, especially when they were never found in homologs (**Figure 3B**). Any structural annotation containing an in-frame stop codon or a frameshift (i.e. any gap in coding sequence that was not an intron but that changed the translation phase), lacking a start codon or a terminal stop codon, or presenting an unexpected splicing site (different from the GT-AG and GC-AG donor/acceptor canonical splicing sites) was called ‘non-canonical’. This careful inspection was facilitated by viewing the sequence annotations with the Artemis editor (Carver et al., 2012).

**Figure 3.**
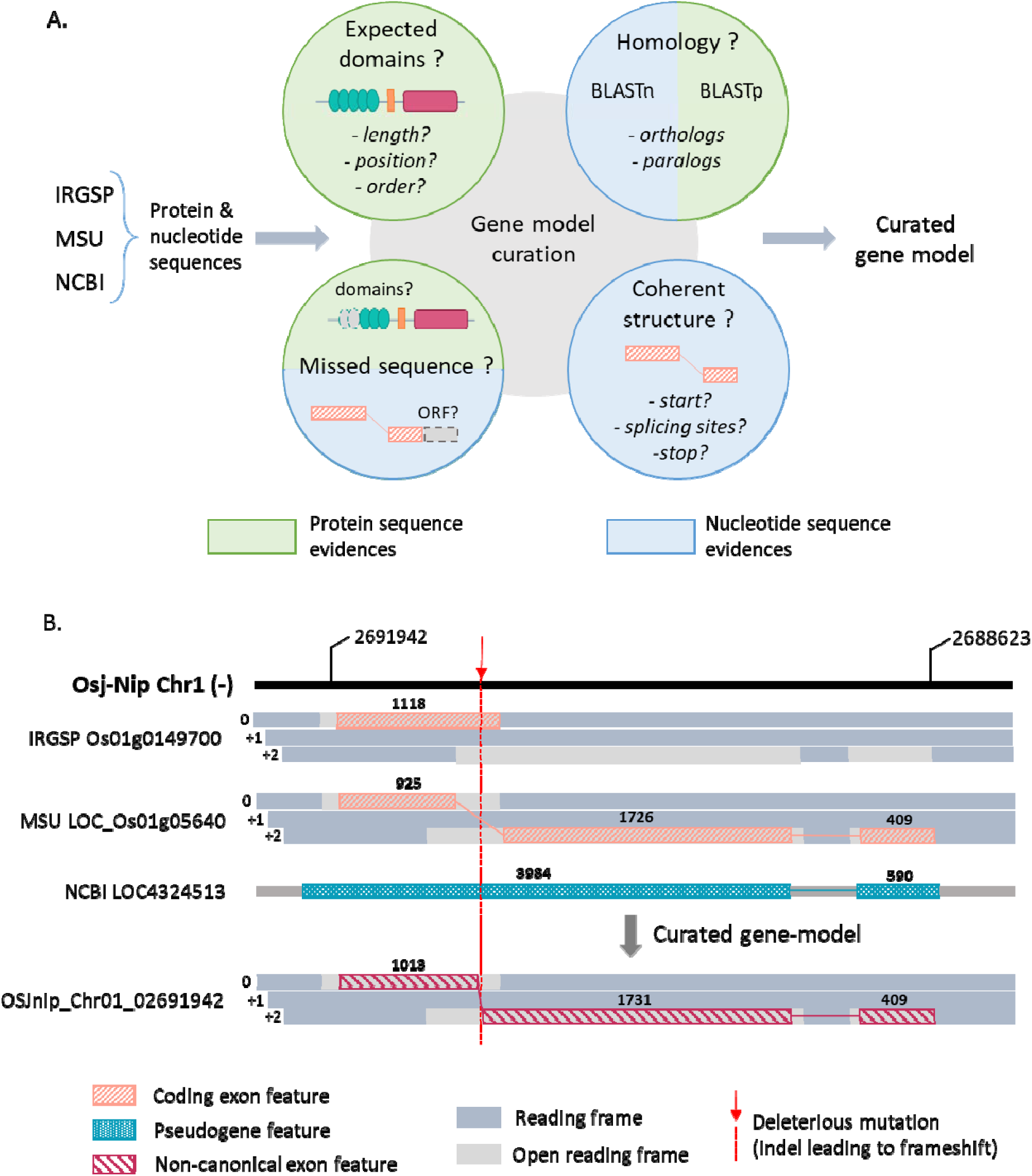
Manual curation of LRR-CR gene models strategy and example of annotation inconsistencies. **(A)** Schematic representation of the strategy used to curate Nipponabre LRR-CR gene models. An initial gene model was selected from the three public annotations. This gene model was gradually modified based on protein and nucleotide sequence evidence. The curated model was then classified as canonical or non-canonical. **(B)** Schematic representation of an example of inconsistency between gene models from publicly available annotations and how the curation was done. The gene is an LRR-RLK located on chromosome 1 of the Nipponbare genome. Numbers above boxes give the feature length. In this example, an indel mutation caused a frameshift in the first exon of the gene. The IRGSP annotation retrieved the first part of the coding sequence stopping at the first stop codon on frame 0. The MSU annotation retrieved a longer coding sequence, but sidestep the indel mutation by introducing a ‘dubious’ intron in order to reach the ORF on the +2 frame. This ‘dubious’ intron was abnormally short and contained a sequence highly homologous to the coding sequence in other paralogous gene copies. The NCBI annotation gave a pseudogene feature, i.e. a feature from which a protein sequence could not be deduced: the cDNA sequence is available but would not allow protein translation as it would be in the wrong reading frame after the mutation. The curation took advantage of the three annotations. It retried a cDNA sequence that overlapped the complete former coding sequence in the two successive correct reading frames via identification of the indel mutation. The identification of the indel mutation was clearcut since the gene was tagged as ‘non-canonical’ with the presence of a frameshift, but it allowed a complete protein sequence to be deduced and used for sequence comparison and alignment.

In a last step, we looked for LRR-containing sequences that would have been missed by the three publicly available annotations. The genome was thus split into 1 kb segments with overlapping 100 bp borders, translated into amino acid sequences in the six reading frames, and domains (LRR, kinase, NB-ARC, etc.) were searched with hmmsearch. The overall results were concatenated and filtered for redundancies. Domains falling inside annotated exons or isolated from other LRR motifs (within a 5 kb range) were discarded. Regions of interest (with at least three LRR motifs in tandem) were compared to other plant genomes using BLAST to screen for the potential presence of a gene model in the considered region.

The final set of manually validated LRR-CR loci on the Nipponbare genome consisted of 1,052 genes (351 LRR-RLKs, 147 LRR-RLPs, 495 NLRs and 59 UCs) (**Table 2** and **Supplemental Data Set 1**). Among these 1,052 genes, eight (1 RLK, 3 RLP and 4 UC) were located at loci for which none of the three publicly available annotations detected a gene. The newly identified LRR-RLK was a canonical full-length sequence on the forward strand of chromosome 2 (from 6,831,702 to 6,834,761). Note that this sequence is actually present in GenBank under the accession number EAZ22278.1, but is located on the reverse strand in a non-coding region of the Os02g0222500 gene. The seven other are non-canonical truncated genes. In addition, for six of these 1,052 validated genes, the LRRprofiler pipeline did not detect any further LRR motifs in the predicted protein. LRR motifs were initially detected for these six genes, but at the threshold limit when using HMM profiles built on the basis of the initial dataset (for details see LRRprofiler pipeline section in the Material and Methods). When using the slightly different HMM profiles obtained with the final dataset, the same LRR motifs were no longer detected as they did not surpass the threshold. However, a careful manual inspection showed that the LRR domain was present but contained divergent LRR motifs, thereby complicating automatic detection. Consequently, these genes were kept and classified according to the presence of the other domains (kinase or NB-ARC). These eleven genes included two LRR-RLKs and nine NLRs.

**Table 2.**
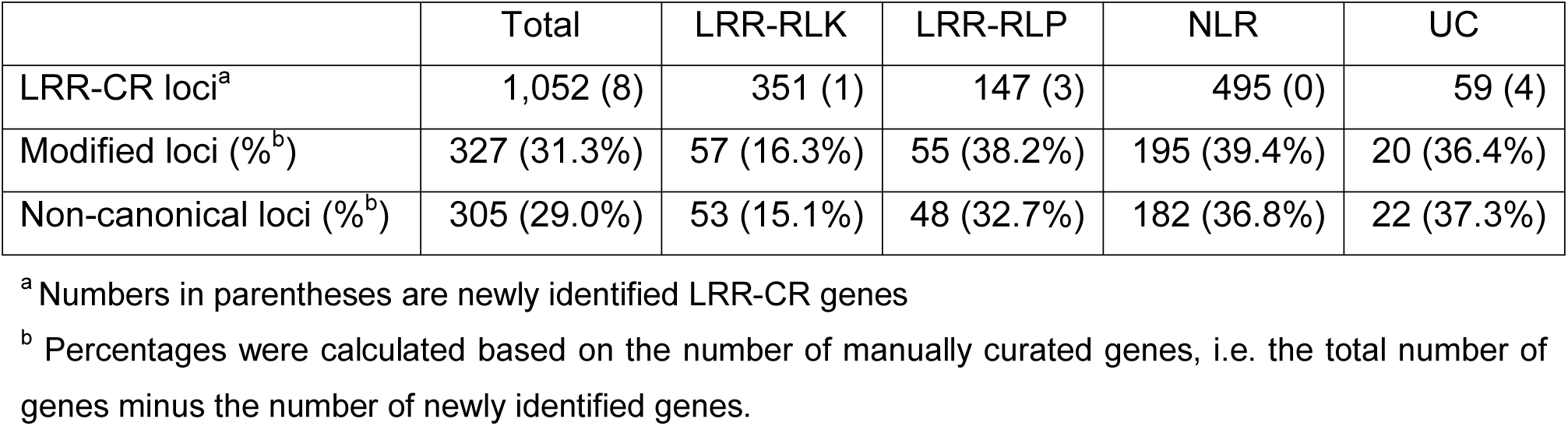
Number of LRR-CR proteins in the predicted proteomes from our curated annotations for the Nipponbare rice reference genome. Sequences were identified and classified into subfamilies using the LRRprofiler pipeline.

|  | Total | LRR-RLK | LRR-RLP | NLR | UC |
| --- | --- | --- | --- | --- | --- |
| LRR-CR loci<sup>a</sup> | 1,052 (8) | 351 (1) | 147 (3) | 495 (0) | 59 (4) |
| Modified loci (%<sup>b</sup>) | 327 (31.3%) | 57 (16.3%) | 55 (38.2%) | 195 (39.4%) | 20 (36.4%) |
| Non-canonical loci (%<sup>b</sup>) | 305 (29.0%) | 53 (15.1%) | 48 (32.7%) | 182 (36.8%) | 22 (37.3%) |

<sup>a</sup> Numbers in parentheses are newly identified LRR-CR genes
<sup>b</sup> Percentages were calculated based on the number of manually curated genes, i.e. the total number of genes minus the number of newly identified genes.

Among these 1,052 LRR-CR genes, 327 (195 NLR, 57 LRR-RLK, 55 LRR-RLP and 20 UC) were manually modified because none of the three publicly available annotations had a satisfactory gene model based on the previously defined criteria (**Figure 3, Supplemental Data Set 1**). The overall proportion of modified loci was 31.3% (327/1,052) and markedly varied according to the considered gene subfamily. Only 16.3% of LRR-RLK loci were modified, while 38.2% of the LRR-RLP loci and 39.4% of the NLR loci were modified (**Table 2**).

Among these 1,052 LRR-CR genes, 305 (29.0%) were non-canonical (**Table 2**). More precisely, 273 genes were both non-canonical and modified, representing 83.5% of the total modified loci (273 over 327) and 89.5% of the non-canonical loci (273 over 305) (**Supplemental Table 1**). The remaining 32 non-canonical genes were either unreported by any of the annotations (7), or reported by the NCBI as pseudogene or gene models having putative errors in the genomic sequence due to differences with respect to the RNAseq data (25).

The different gene subfamilies did not contain the same proportion of non-canonical gene models (**Table 2**). Very similar to what was observed regarding the proportion of modified gene models according to the gene subfamily, non-canonical gene models concerned only 15.1% of the LRR-RLK, compared to 32.7% and 36.8% of the LRR-RLP and NLR, respectively.

One way to assess the relevance of our expert LRR-CR annotations is to compare the number of functional domains (TM, NB-ARC, kinase, and LRRs) found in LRR-CR proteins derived from the reference annotations to the number of functional domains found in the proteins derived from our expert annotation (**Supplemental Figure 2**). These comparisons revealed that quite a few more LRR-CR domains were found in our manual annotation as compared to the publicly available annotations. For example, when compared to the IRGSP annotation, our expert annotation highlighted 29% more TM, 42% more NB-ARC, 33% more kinase and 20% more LRR motifs.

### Comparison of LRR-CR loci between the Nipponbare and Kitaake rice cultivars

Kitaake is another *O. sativa* ssp. *japonica* variety for which a complete genomic sequence is available (Jain et al., 2019). In order to compare the LRR-CR repertoire between Nipponbare and Kitaake and limit the need for manual curation in the re-annotation of this closely related rice cultivar, we have developed a strategy to transfer our expert annotations from the Nipponbare to the Kitaake genome.

The strategy summarized in **Figure 4** starts by identifying Kitaake genome regions that are homologous to Nipponbare LRR-CR sequences. Then it successively takes into account three levels of annotation transfer, depending mostly on the level of sequence identity of each considered region with the LRR-CR gene that identified it. At each locus, our strategy strives to retrieve the most probable gene model with the idea that, if possible, it should be canonical. At the end of the process, LRR-CR gene models that are found to be non-canonical or having a dubious protein structure in Kitaake are manually checked and corrected if needed. At this step, the transfer allowed us to identify 1046 genes.

**Figure 4:**
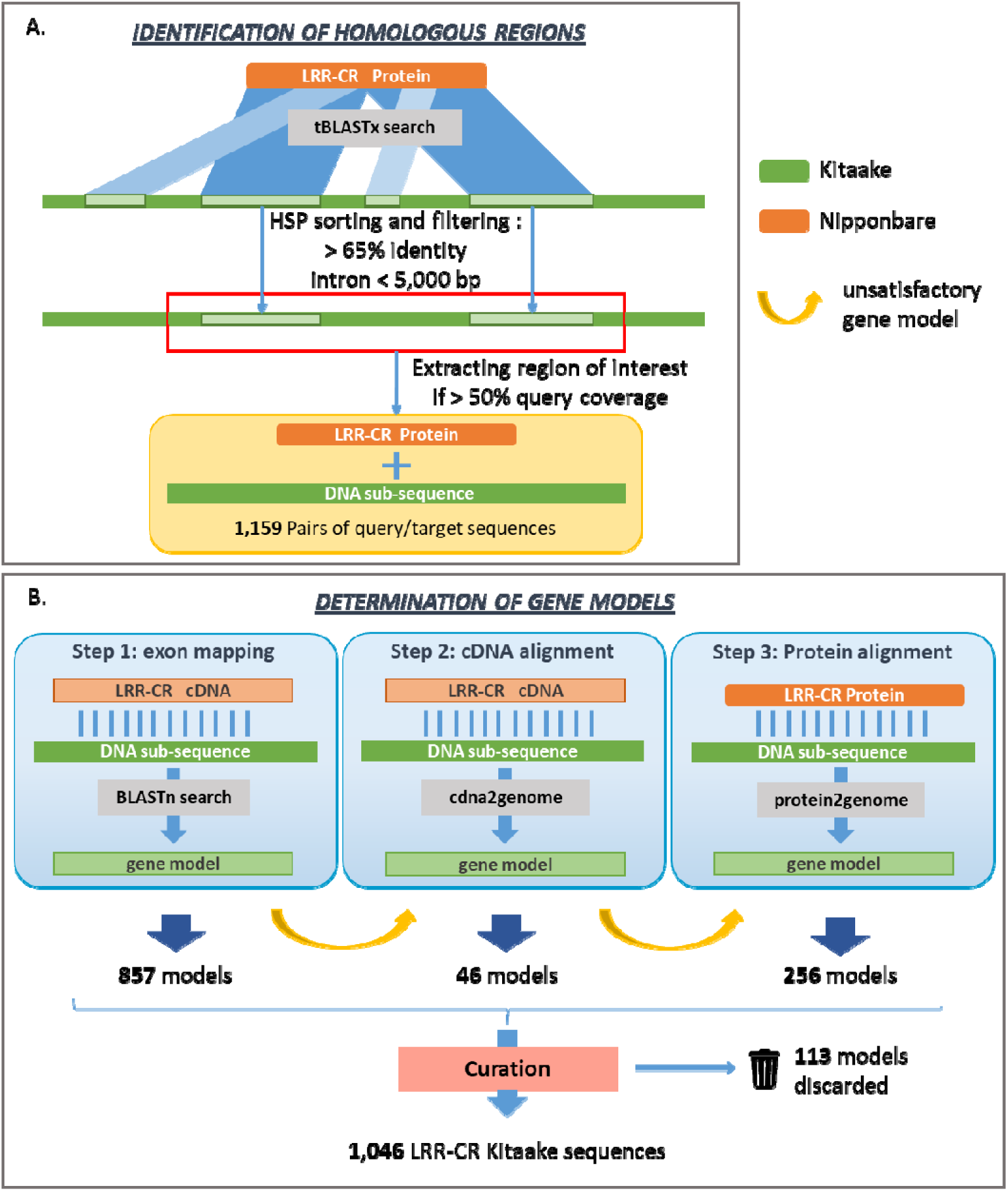
Schematic representation of the annotation transfer strategy between closely related genomes. **(A)** Identification of LRR-CR homologous regions. Nipponbare LRR-CR proteins were used to search regions of interest in the Kitaake genome using tBLASTx. BLAST hits with over 65% identity were ranked in the LRR-CR query protein sequence order and used to define the region boundaries. If the filtered BLAST hits within a Kitaake region covered more than 50% of the query LRR-CR sequence, then the Kitaake sequence of this region was extracted and linked to the Nipponbare LRR-CR query protein. **(B)** Determination of gene models. The process strives to give a gene model for each region of interest identified in the Kitaake genome. The annotation is attempted in three consecutive steps. If the model from one step is unsatisfactory, i.e. gives an alignment of poor quality with the Nipponbare query protein, the process goes to the next step for this region. At the end of the third step, gene models that remained unsatisfactory were manually checked. This process allowed us to annotate 1,046 genes in the Kitaake genome.

As carried out for Nipponbare, the Kitaake genome was finally scanned with LRR HMM profiles for new LRR-CR identifications. This procedure allowed us to annotate 18 additional genes, thereby leading to a total of 1,064 LRR-CR genes in the Kitaake genome.

The LRRprofiler pipeline was used on the 1,064 predicted Kitaake proteins and allowed us to detect LRR in 1,054 of them, 998 were further classified in a LRR-CR subfamily and 56 remained in the UC group. Automatic detection of LRR failed for 10 genes. As carried out for Nipponbare, at this step, manual validation of the protein annotations confirmed the presence of an LRR domain of the expected size and located at the expected positions. These 10 genes were therefore kept in the final dataset. The gene subfamilies for these 10 loci were determined based on other functional domains and a homology search against other LRR-CR protein sequences. Finally, the LRR-CR gene set from Kitaake was composed of 360 LRR-RLKs, 140 LRR-RLPs, 508 NLRs and 56 UCs (**Figure 5** and **Supplemental Data Set 2**).

**Figure 5.**
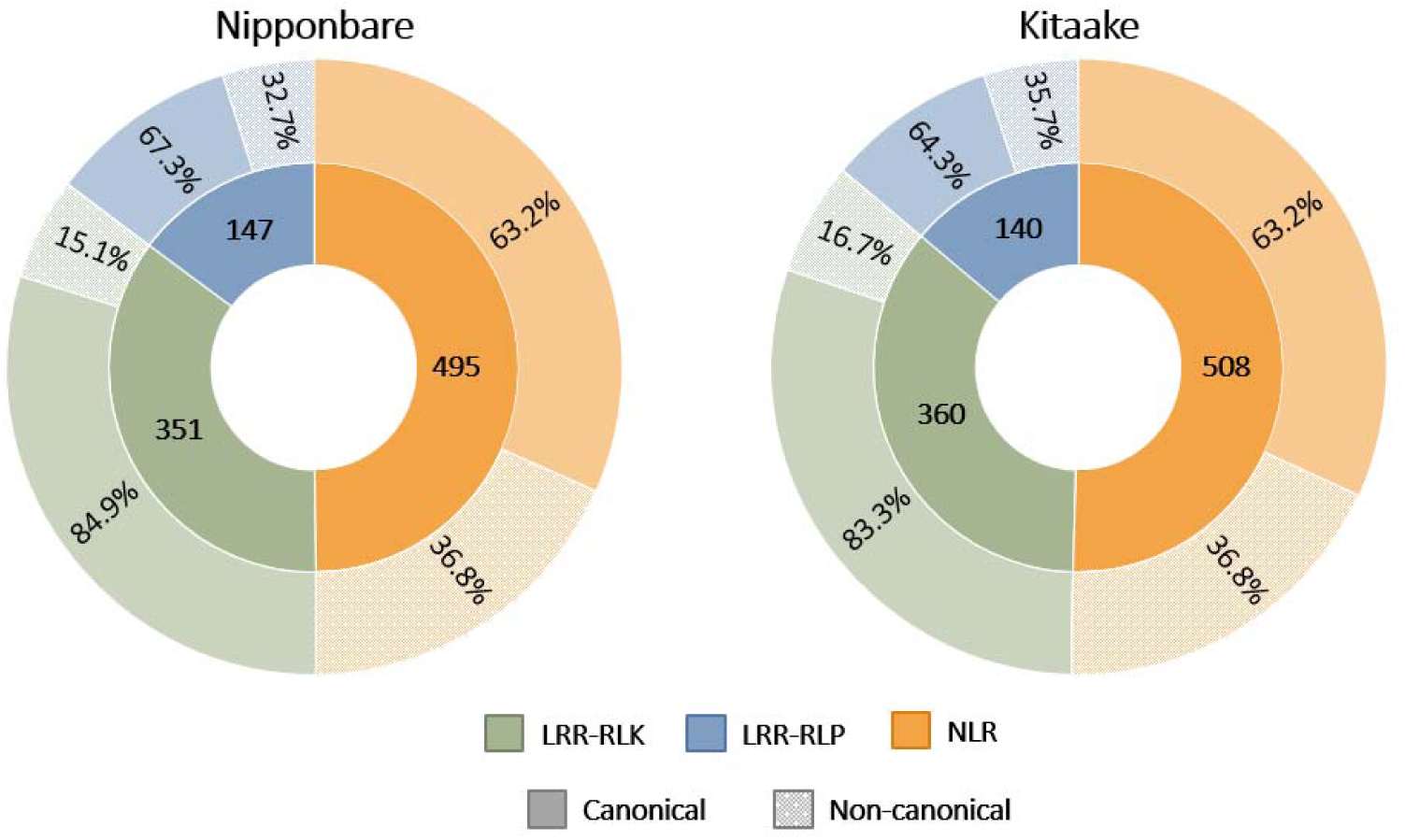
Proportion of canonical and non-canonical loci per gene subfamily in our Nipponbare and Kitaake expert annotations. Percentages were calculated per gene subfamily. The inner circle provides the number of loci per family, with a different colour for each. The outer circle shows a lighter (respectively darker) version of the loci family colour to represent the fraction of the non-canonical (respectively canonical) members within this gene family.

We then tagged all of these Kitaake LRR-CR gene models as either canonical or non-canonical. We obtained 744 (69.9%) canonical genes and 320 (30.1%) non-canonical genes. The proportions of canonical and non-canonical genes per subfamily for Kitaake were very similar to those obtained for Nipponbare (**Figure 5**).

All LRR-CR loci annotation and sequence data for the Nipponbare and Kitaake genomes can be viewed and downloaded on the dedicated website (https://rice-genome-hub.southgreen.fr/content/geloc).

### Comparison of LRR-CR alleles between Nipponbare and Kitaake

Nipponbare and Kitaake are two varieties of the same subspecies, i.e. *Oryza sativa* ssp. *japonica*. As such, for the majority of the genes found in Nipponbare, an allele (i.e. a version of the same gene located at the same chromosomal location) was expected to be found.

By using the Synmap program (Lyons and Freeling, 2008), we identified 999 allelic pairs (representing 90.0% of the total number of loci) between Nipponbare and Kitaake (**Supplemental Data Set 3**). In addition, we noticed that for three NLR gene pairs located close to each other on chromosome 9 in the Nipponbare genome, three consecutive genes of the Kitaake chromosome 3 were found with 100% identity with regard to their predicted coding sequence. The intergenic sequences of these two regions also had a high level of identity (99% over 40.5 kb), suggesting that these three genes are located in a translocated region of the genome. Among these 1,002 pairs (999 allelic plus 3 translocated), 688 (68.7%) were pairs of canonical gene models, 267 (26.6%) were pairs of non-canonical gene models, and 47 (4.7%) were pairs of genes found to be canonical only in one of the two cultivars. Interestingly, 83.3% of the LRR-RLK pairs were canonical in both cultivars, compared to only 63.9% of the LRR-RLP pairs and 60.5% of the NLR pairs (**Table 3**).

**Table 3.**
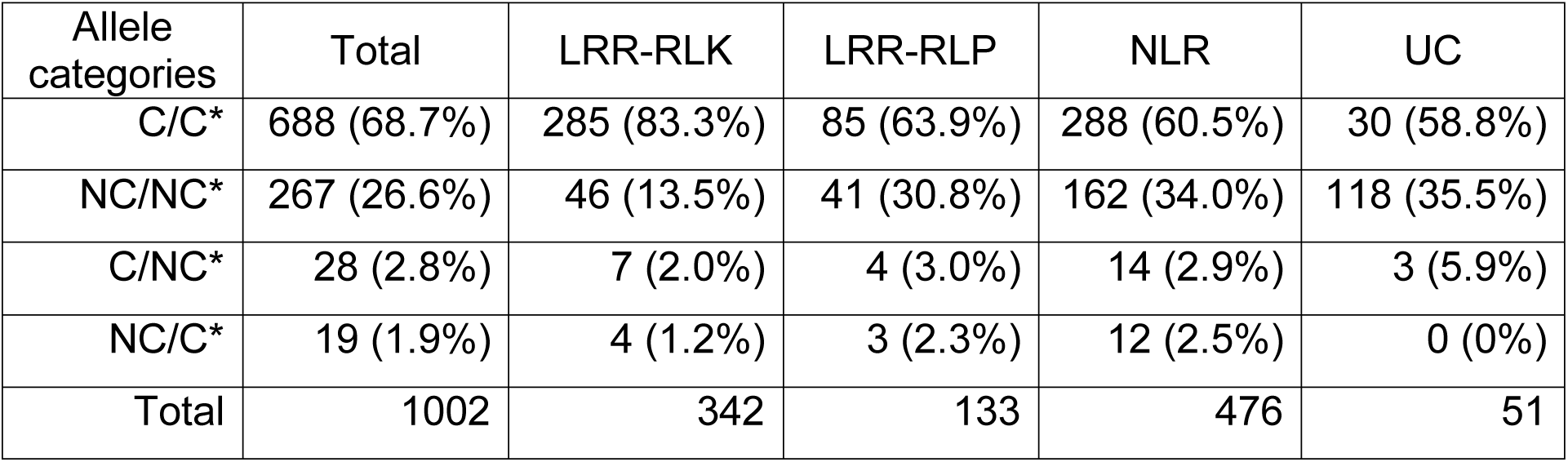
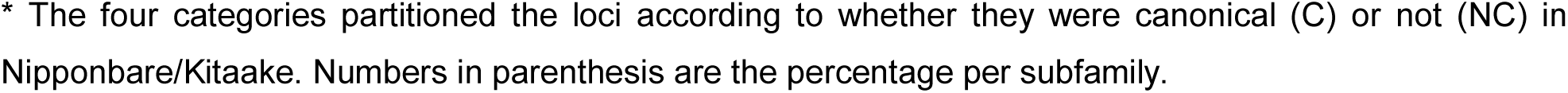
Number of allelic pairs between Nipponbare and Kitaake cultivars according to categories and subfamilies.

| Allele categories | Total | LRR-RLK | LRR-RLP | NLR | UC |
| --- | --- | --- | --- | --- | --- |
| C/C* | 688 (68.7%) | 285 (83.3%) | 85 (63.9%) | 288 (60.5%) | 30 (58.8%) |
| NC/NC* | 267 (26.6%) | 46 (13.5%) | 41 (30.8%) | 162 (34.0%) | 118 (35.5%) |
| C/NC* | 28 (2.8%) | 7 (2.0%) | 4 (3.0%) | 14 (2.9%) | 3 (5.9%) |
| NC/C* | 19 (1.9%) | 4 (1.2%) | 3 (2.3%) | 12 (2.5%) | 0 (0%) |
| Total | 1002 | 342 | 133 | 476 | 51 |

\* The four categories partitioned the loci according to whether they were canonical (C) or not (NC) in Nipponbare/Kitaake. Numbers in parenthesis are the percentage per subfamily.

The remaining 111 genes were genotype-specific: 49 were present only in Nipponbare and 62 only in Kitaake, of which 31 (63.3%) and 37 (59.7%), respectively, were canonical. These genes were not evenly distributed on the genomes. Some of them were located in entirely different regions in Nipponbare and Kitaake, in particular in chromosomes 2, 11 and 12 where entire clusters were genotype-specific (**Figure 6A**). Other genotype-specific genes were dispersed in regions containing conserved allelic pairs in either Nipponbare or Kitaake (**Figure 6B**). Note that chromosome 11, which also contained about a fifth of all LRR-CR genes, hosted 44 of the 111 (40.4%) cultivar-specific loci.

**Figure 6.**
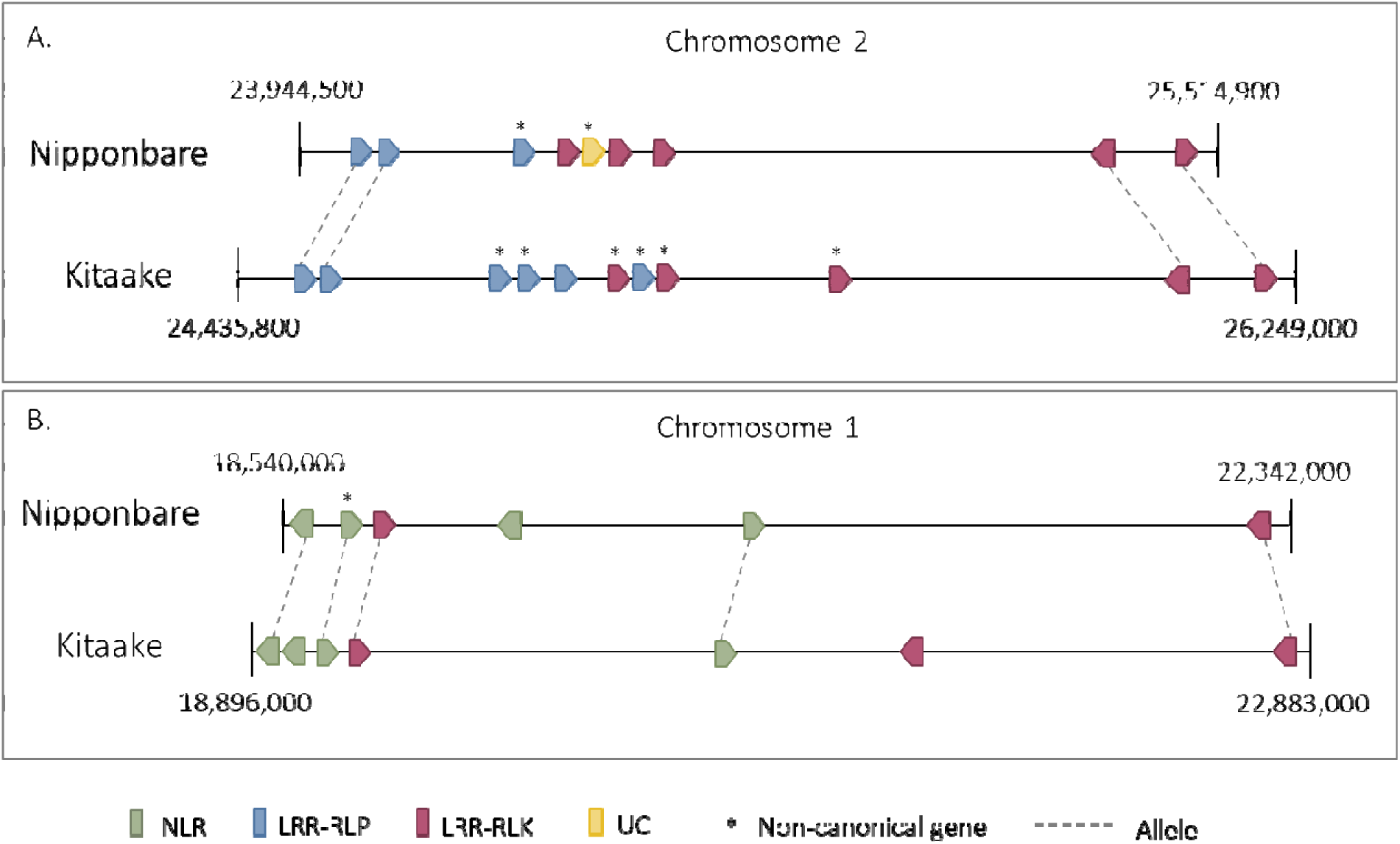
Schematic representation of two large loci on chromosomes 1 and 2 containing cultivar-specific LRR-CR genes. **(A)** Representation of an unconserved cluster between Nipponbare and Kitaake on chromosome 2. Five and seven genes in Nipponbare and Kittake, respectively, were cultivar specific. The unconserved region was framed by four conserved genes, i.e. two LRR-RLPs and two LRR-RLKs. **(B)** Representation of a conserved region between Nipponbare and Kitaake on chromosome 1 hosting cultivar-specific genes. The Nipponbare region hosted a cultivar-specific NLR while the Kitaake corresponding region hosted two cultivar-specific genes, i.e. an NLR and an LRR-RLK.

We checked one specific feature of the sequenced Kitaake genome (KitaakeX), i.e. the fact that it contains two *XA21* transgenes (Song et al., 1995; Jain et al., 2019). We identified these two transgenes at positions 28,161,378 and 28,165,947 on chromosome 6 with no corresponding allele in Nipponbare, in accordance with published data.

For each of the 999 pairs of LRR-CR alleles (plus the three translocated pairs), the fraction of exact matches along the cDNA pairwise alignment (i.e. their percentage of identity) was computed. This cDNA identity was about 99.0% on average. The highest identity rate (99.6%) was obtained for alleles belonging to the LRR-RLK subfamily, followed by the NLR (98.7%) and LRR-RLP (98.0%). Non-canonical conserved gene pairs (NC/NC category in **Table 3**) had, on average, a lower identity level (98.7%) than the conserved canonical gene pairs (99.4%). The lowest level of sequence identity (93.4%) was noted between gene pairs with one cultivar having a canonical form and the other a non-canonical form (categories C/NC and NC/C in **Table 3**). Only 17 pairs of alleles shared less than 80% cDNA identity due to both deletions (up to 1.7 kb) and high sequence divergence (**Supplemental Table 2**).

### Impact of reannotations on LRR motif numbers in alleles

To obtain a precise annotation of LRR motifs in each protein, we used the LRRprofiler pipeline on proteomes predicted by the publicly available annotations on one hand, and by our manually curated annotations on the other, for both cultivars. A total of 999 pairs of alleles between Nipponbare and Kitaake were identified from our manual annotation (plus three translocated pairs, see above). The same procedure was applied to retrieve pairs of alleles of publicly available annotations. Then the number of LRR motifs predicted in sequences was compared between Nipponbare and Kitaake alleles.

We observed a mean difference in LRR number per protein of 3.58 when comparing the publicly available annotations (IRGSP for Nipponbare and the only one that exists for Kitaake) (**Figure 7, Supplemental Figure 3**). This difference fell to 0.6 when our re-annotated data were compared. Using our curated annotations hence led to LRR number predictions that were much more consistent between Nipponbare and Kitaake alleles and this trend was observed for all LRR-CR gene subfamilies. Moreover, the mean difference in LRR number still varied between LRR-CR gene subfamilies, with greater conservation of LRR motif numbers between LRR-RLK and LRR-RLP alleles than between NLR alleles.

**Figure 7.**
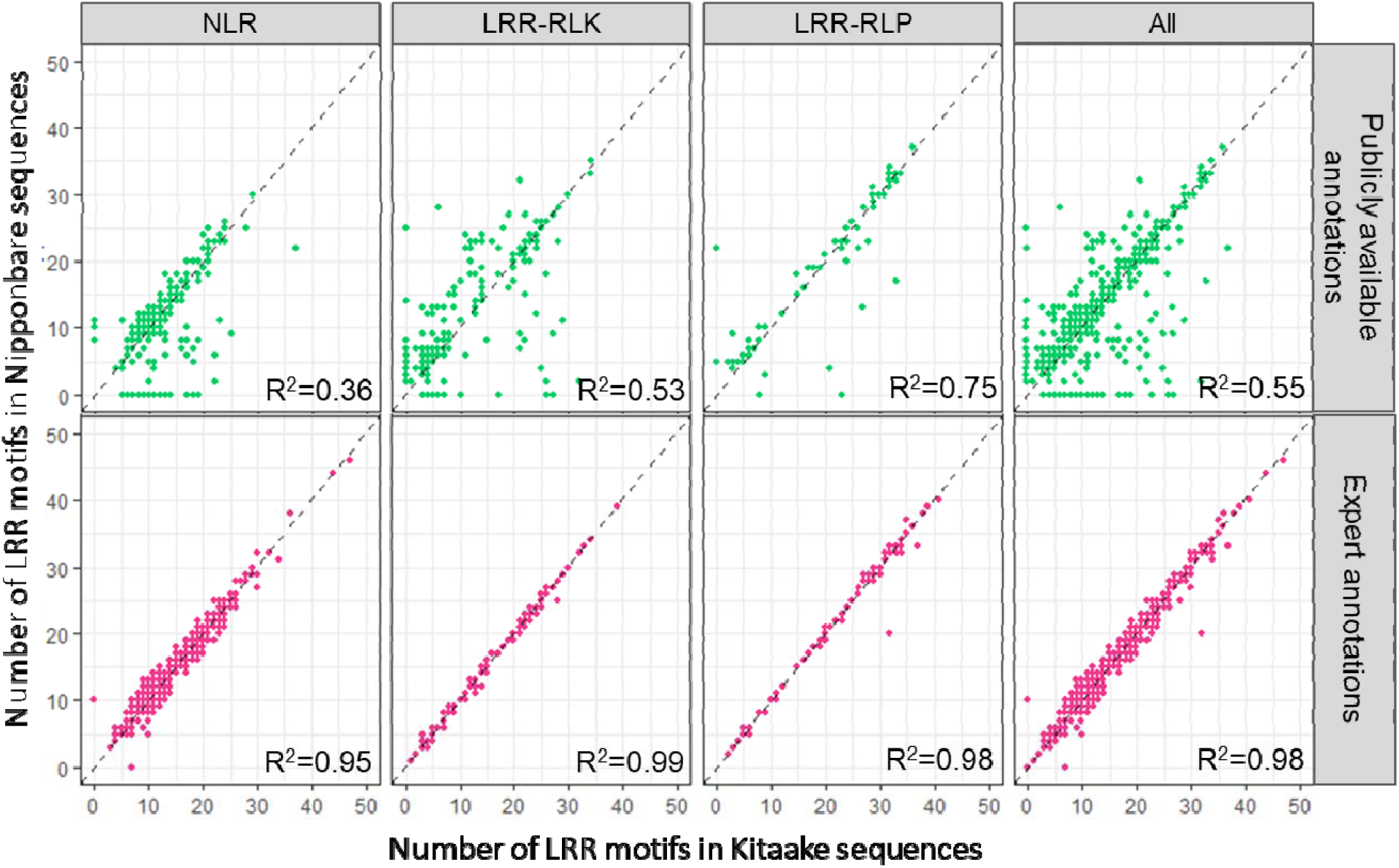
Comparison of LRR motif numbers between Nipponbare and Kitaake LRR-CR alleles, according to the annotations used. At the top, in green, comparison between two publicly available annotations of Nipponbare and Kitaake using IRGSP reference data for Nipponbare; at the bottom, in pink, comparison between our Nipponbare and Kitaake expert annotations.

## Discussion

In recent years, in the wake of the gigantic amount of genome sequenced data available, it was exciting to undertake evolutionary studies of gene families. We have been part of this collective enthusiasm, and like many others, have based our research conclusions on perfectible versions of automatic structural and functional gene annotations (Fischer et al., 2016; Dufayard et al., 2017). Although previous genome-wide phylogenetic approaches on LRR-CR gene families enhanced our knowledge on their evolution, they almost never included a data curation step. Indeed, manual re-annotation of gene families is a laborious time-consuming task, especially when dealing with large complex gene families such as LRR-CR, and even more so when dealing with many plant genomes. Despite the fact that automatic annotation tools are continuously improving and remain indispensable at the genome level, human expertise is clearly still needed to achieve an annotation accuracy level suitable for finer and deeper analyses. Today we have finally undertaken this re-annotation work anyway because we are convinced that these curated data are required to produce new reliable results on the evolution of these gene families, especially in the current pangenomic era. Here we describe a new so-called ‘comprehensive’ annotation strategy. We hope that this new annotation process will get its place in addition to structural and functional annotations in use thus far.

### Automatic annotations give inconsistent gene models on complex multigenic families

Our comparison of the three publicly available annotations for the Nipponbare rice reference genome showed major discrepancies regarding the total number of LRR-CR genes, the number of LRR-CR assigned to each subfamily, and the gene models (**Table 1, Figure 2** and **Supplemental Figure 1**). These differences were greater for LRR-CR genes compared to other genes such as TF. Automatic annotation pipelines appeared to be suitable overall for many gene families, but they led to a large proportion of inconsistent gene models when annotating complex multigenic families like LRR-CR. The annotation of fast-evolving multigene families is especially challenging for automatic approaches because a high duplication rate is often accompanied by a loss of function process (pseudogeneization) for many copies due, for instance, to mutations like nucleotide substitutions, which may introduce premature stop codons or indels, in turn generating frameshifts. This can lead to the presence of several gene copies sharing high sequence similarity even though some of them may contain nonsense mutations, frameshifts or be truncated. The annotation of gene copies harbouring nonsense mutations is problematic compared to the initial unaltered copy. Some pipelines will be able to detect the entire coding phase but will introduce false introns to sidestep stop codons or frameshifts in order to retrieve a putatively translatable CDS (**Figure 3**). We noticed that the MSU automatic annotation tended to sidestep nonsense and frameshift mutations by introducing short introns. Such errors were observed previously by Meyers when re-annotating the *A. thaliana* NLR gene family (Meyers et al., 2003). Two arguments strengthen the assertion that such introns are false: (i) such introns are never found in more than one copy whereas the intron positions are known to be well preserved between closely related copies, and (ii) sequence comparisons performed against recent paralogues (or orthologues from close relative species) have shown that the sequences of these wrongly annotated introns are always clearly homologous to coding sequences in other gene copies. Among the intron gain mechanisms, intronization (i.e. the process by which an exonic sequence is changed into an intron by mutation accumulation) is a complex process that is not yet very well documented or understood (Yenerall and Zhou, 2012; Roy, 2016). If it really occurs in genomes, it implies that a sufficiently long enough period of time must have passed for these mutations to occur and generate novel splicing sites. It is thus very unlikely that so many new introns arose in such a short period of time, as revealed by the low level of divergence between these genes and their paralogue and/or orthologue counterparts. Other annotation pipelines, such as that of the IRGSP consortium, are more conservative in the sense that they give gene models with a more biologically meaningful expected structure, e.g. truncated proteins, in accordance with the presence of the first premature stop codons (either in-frame or caused by a frameshift, **Figure 3**). This conservative choice could likely explain why we found more sequences classified as LRR-RLP from the IRGSP annotation than from the two others (**Table 1**). Indeed, any LRR-RLK with a premature stop codon somewhere before the kinase domain would be considered as LRR-RLP (**Figure 1, Figure 3**). The annotation inconsistencies that we pinpointed here were observed for LRR-CR genes and did not question the overall quality of the three available rice genome annotations. They highlighted the limits of automatic annotation pipelines to annotate such complex multigene families and the consequences that given pipeline decision rules may have when drawing evolutionary conclusions.

The comparisons of the different gene models proposed by the three Nipponbare annotations led us to undertake a manual curation of the LRR-CR gene family. We are not the first to get involved in this painstaking but necessary work. Several high-quality studies have been based on re-annotated data, particularly in *A*.*thaliana* (Meyers et al., 2003; Van de Weyer et al., 2019). Expert annotations could also of course contain errors, but expert curation limits their number. Opting exclusively for automated annotation should be avoided or otherwise operators should be aware that these annotations may contain errors induced by gene family specificities. These biases must be known and understood to avoid drawing misleading conclusions.

### The expert Nipponbare rice genome contains more than 1,000 LRR-CR loci of which 30% have a non-canonical gene model

We curated LRR-CR loci in the reference Nipponbare rice genome by first comparing the three publicly available annotations at each locus: IRGSP, MSU and NCBI. Our aim was to retrieve LRR-CR genes in their entirety, while taking the coding sequences as they probably were before mutation accumulation into account. We obtained evidence that the sequence portions we included in our new gene models were not random genomic sequences but instead parts of the original gene CDS, as shown by the recovery of protein domains belonging to LRR-CR genes (kinase, NB-ARC, TM. **Supplemental Figure 2**). When a gene had a nonsense mutation (in-frame stop codon or frameshift), an unexpected splicing site, or no terminal stop or start codon, we tagged it as non-canonical. This canonical vs. non-canonical classification was based solely on features observed in gene models and did not imply any judgements on gene functionality. Genes tagged as non-canonical spanned a wide variety of cases, some of which could very likely not be translated into a functional protein while others may have had a function. As a first example, mutations inducing a premature stop codon could lead to a shorter protein that might sometimes perform the same function. Yet in many other cases shorter proteins might not perform the same function, if able to perform any function at all. Another example concerns loss of the expected stop codon. When screening a sequence, a stop codon will eventually be encountered, but determining the functional consequences of this additional amino acid stretch would be impossible *in silico*. The same holds when the start codon is lost. Determining the criteria according to which an alternative start codon (if any) may become the new start codon, is a hazardous task. These few examples highlight the extent to which sorting out different functional scenarios is challenging. Moreover, mRNA molecules may play a regulating role, even if they cannot be translated as such, thereby justifying the need to annotate them. These reflexions led us to voluntarily disregard such interpretations in our re-annotation process.

We observed that a third of the LRR-CR genes were non-canonical overall, but their proportion varied according to the gene subfamily (**Table 2**). A lower proportion of LRR-RLK genes were non-canonical (15%), compared to LRR-RLP (32%) and NLR (37%). The LRR-RLK subfamily could be divided into 15-20 subgroups based on phylogenetic study findings, and the duplication rate was shown to be quite variable according to the subgroup considered (Tang et al., 2010; Fischer et al., 2016). Some subgroups, whose genes have been described to be mostly involved in developmental processes, have had a more stable copy number over the course of angiosperm evolution (Fischer et al., 2016). These genes are less prone to duplication, and thus less likely to generate copies accumulating nonsense mutations, thereby lowering the proportion of non-canonical genes when the entire LRR-RLK subfamily is considered. The higher proportion of non-canonical genes obtained for NLR and LRR-RLP suggests that these subfamilies generally have higher birth and death rates. A quarter of the LRR-CR genes required manual curation and were non-canonical (representing 83.5% of the curated loci and 89.5% of the non-canonical loci (**Supplemental table 1**)). In fact, manual curation was conducted mainly when none of the three annotations gave satisfactory gene models (such as the example presented in **Figure 3B**). The high correlation between non-canonical and curated loci was thus likely due to the presence of mutations introducing ambiguities, which are overcome to different extents by the three annotation pipelines.

We stress that this categorization of loci into a canonical or non-canonical models could be impacted by the genomic sequence quality. Errors in the reference genome sequence could introduce errors in the gene models. In our curated dataset, 27 non-canonical genes were tagged in NCBI data as harbouring a difference when considering the reference genome sequence and the RNAseq data. The mutations jeopardizing the expected gene structure corresponded exactly to the positions where inconsistencies had been highlighted between the genomic and RNAseq sequences. Without RNAseq data, genome sequence errors cannot be differentiated from real mutations. Redundancies in LRR-CR gene sequences can give rise to ambiguities during both genome sequence assembly and expression data mapping, thus leading to errors (Torresen et al., 2019). Only specific re-sequencing could resolve those potential inconsistencies. In rice, this concerns only 2.6% of the genes in the current state of the data presented in this study.

### LRR-CR repertoire in Kitaake and comparison with Nipponbare

We propose a modular strategy to transfer our manually curated annotations to other rice genomes. We applied this strategy to annotate LRR-CR genes from the genome of the Kitaake cultivar, which also belongs to the *O. sativa* Japonica subspecies. A comparison of the Nipponbare and Kitaake LRR-CR repertoires revealed an equivalent number of loci. The distributions of LRR-CR loci per gene subfamily, chromosome and category (canonical or not) were also consistent between these two cultivars (**Figure 5**). In the Nipponbare genome, eight new LRR-CRs (1 LRR-RLK, 3 LRR-RLPs, and 4 UCs) were identified. These genes had not been previously annotated in any of the three publicly available annotations. In Kitaake, the same strategy enabled us to identify 114 new LRR-CR genes (48 of which were canonical): 17 LRR-RLKs, 24 LRR-RLPs, 50 NLRs and 23 UCs. The higher number of newly identified genes in Kitaake (114 vs 8) suggested that annotation inaccuracies had a greater impact on recently sequenced genomes that have not benefited from as much annotation investment as reference genomes.

A comparison of the LRR motif number for all allelic pairs between Nipponbare and Kitaake revealed a much greater difference in LRR number between alleles for publicly available annotations (ranging from 2.68 to 3.58) in comparison to our manually curated annotations (0.58) when the three subfamilies were all considered (**Figure 7** and **Supplemental Figure 3**). When publicly available annotations are considered, some rare allelic pairs harboring a very different number of LRR may be truly different functional alleles. For instance, between two LRR-RLP alleles, one may contain a premature stop-codon leading to the loss of a few motifs, but it may still have a biological function. It is important to identify such a pair. In our expert annotation, alleles may share an identical number of LRR but such allelic pairs would be clearly identifiable because one of the alleles would be tagged as non-canonical while the other would be canonical. Moreover, in non-canonical alleles, the causal mutation, its position and impact on the gene (i.e. frameshift or premature stop-codon) could be identified.

The difference in LRR motif number observed between allele pairs was greater for NLR than for the other subfamilies (**Figure 7**). This might be explained by the fact that NLR motifs are more variable and hence harder to detect (Ng et al., 2011), which could lead to apparent variations in the number of motifs in the two alleles. LRR motifs that have been found in NLRs differed from the common “Plant Specific” LRR consensus sequence, and were more irregular in terms of both length and residue conservation (Kajava, 1998; Kuang et al., 2004; Matsushima and Miyashita, 2012; Sela et al., 2012). Although we enhanced the LRR detection accuracy through the development of a new LRR HMM profile for NLR, it is still not exhaustive. This also suggests that the number of LRR motifs varies more in NLR than in other LRR-CR subfamilies.

High sequence similarity was observed between Nipponbare and Kitaake alleles (98.9% identity for cDNA) (**Supplemental Table 2**), which was consistent with previous comparisons (Jain et al., 2019). However, some allelic pairs showed a lower identity level with a more ancient coalescent history between the two genomes. This heterogeneity may have been the consequence of breeding programs from which these varieties were derived. The breeding process involves crosses with more or less closely related genotypes, sometimes from different subspecies, and may generate mosaic genomes (Santos et al., 2019). No allelic pairs were found for 111 genes, i.e. 49 were specific to Nipponbare and 62 to Kitaake. A majority of those genes were located in clusters on chromosomes 2, 11 and 12, which have already been described as containing a large number of LRR-CRs (Zhou et al., 2004; Mizuno et al., 2020)(**Figure 6**). Some clusters have also been shown to be less conserved (Mizuno et al., 2020). More than half of these genes were classified as canonical, so they would generally be more effectively functional.

The methods we developed allowed us to undertake an exhaustive comparison of the LRR-CR repertoire between Nipponbare and Kitaake. Allelic pairs, including those hosting nonsense mutations in either or both genotypes, were described (**Supplemental Data Set 3**). Genotype-specific genes were also identified and localized, again along with information related to the potential presence of nonsense mutations (**Figure 6**). These results were achieved through a combination of an expert annotation and its transfer to a second genotype for which a high quality *de novo* genome assembly was available. Validation of the LRR-CR annotations of Kitaake was not very time consuming compared to the initial work in Nipponbare where each gene was investigated individually. Our study highlighted that investment in long-read sequence technologies would be warranted in order to be able to work with high quality *de novo* assemblies, especially when allelic diversity discovery is targeted.

The tools and curated datasets we generated in this study are available on these websites: https://rice-genome-hub.southgreen.fr/content/geloc (data) and https://github.com/cgottin/LRRprofiler (tools). We believe that evolutionary studies and allele discovery initiatives on LRR-CRs would be more accurate and reliable when using our new manually curated ‘comprehensive’ annotations for these genes. Moreover, we feel that this ‘comprehensive’ annotation approach should be widely adopted by the community in the light of its major potential benefits.

## Material and Methods

### Genomes and annotation files

Reference genomic sequences of Nipponbare (Kawahara et al., 2013) and Kitaake (Jain et al., 2019) *Oryza sativa* ssp. *japonica* cultivars were downloaded from the Rice Annotation Project Database (RAP-DB) website (https://rapdb.dna.affrc.go.jp/) and the Phytozome website (https://phytozome.jgi.doe.gov/pz/portal.html). The general feature format (GFF) and fasta files with coding DNA sequences (CDS) and protein sequences for Nipponbare were downloaded for three different annotation projects: (i) the MSU (v7.0) annotation was downloaded from the Rice Genome Annotation Project FTP server (http://rice.plantbiology.msu.edu/), (ii) the IRGSP annotation files were downloaded from the RAP-DB website (https://rapdb.dna.affrc.go.jp/), and (iii) the NCBI annotation (release 102) annotated by the NCBI Eukaryotic Genome Annotation Pipeline was downloaded from the NCBI website (https://www.ncbi.nlm.nih.gov/). The IRGSP annotation consists of two gene sets (‘genes supported by FL-cDNAs, ESTs or proteins’ and ‘computationally predicted genes*’*) that were concatenated for the analyses (Sakai et al., 2013).

### LRRprofiler implementation

The LRRprofiler pipeline was implemented in two steps (**Supplemental Figure 4**). The first step involved iterative refinement of LRR HMM profiles specific to a gene subfamily (LRR-RLK or NLR) and proteome (inspired by (Ng et al., 2011), **Supplemental Figure 4A**). Only LRR-RLK and NLR were considered for profile refinement because they contain a specific domain (i.e. kinase and NB-ARC domains, respectively), thereby allowing clear identification of the subfamily to which they belong. A set of candidate protein sequences was identified from a given proteome to refine the specific LRR profiles. This set was composed of either LRR-RLKs identified with iTAK (Zheng et al., 2016) or NLRs identified with the PF00931 Pfam NB-ARC profile. A first round of LRR motif detection was performed in either of the candidate protein sets using hmmsearch (HMMER, (Eddy, 2011)) with the SM00370 LRR profile from the SMART database. Motifs of 20 to 26 amino acids in length were extracted, aligned with MAFFT with default parameters and a new profile was built from the alignment using hmmbuilt (HMMER, (Eddy, 2011)). This process was repeated using the HMM LRR profile built at the previous iteration to search again for LRR motifs in the considered protein candidate set. At each iteration, the sum of the amino acid lengths of the detected LRR motifs was calculated. The process stopped when three iterations (not necessarily consecutive) resulted in a decrease of this statistics. Finally, the process retrieved the HMM LRR profile identifying the maximum number of LRR motifs in the candidate protein set.

The second step of the LRRprofiler pipeline consisted of identification of LRR-CR proteins contained in a given proteome, annotation of their functional domains, as well as their classification into a gene subfamily: LRR-RLK, LRR-RLP, NLR or UC (**Supplemental Figure 4B**). Four publicly available LRR HMM profiles from the SMART database, i.e. SM00367 (LRR_CC), SM00368 (LRR_RI), SM00369 (LRR_TYP) and SM00370 (LRR), in addition to the newly built LRR profiles obtained in the first step, were used to detect LRR motifs in the complete considered proteome using hmmsearch. An annotation of LRR domains containing the start and end positions of each LRR motif was output. The annotation of each protein was then supplemented using publicly available profiles for other functional domains: TIR (PF01582), Malectin (PF11721), Malectin-like (PF12819), RPW8 (PF05659), Cys-Pairs (Dievart and Clark, 2003; Dufayard et al., 2017), F-box (PF00646) and FBD (PF08387). NB-ARC and kinase domain annotations were retrieved from the first step, while transmembrane domains (TM) were detected with TMHMM (v2.0c) with default parameters (Sonnhammer et al., 1998). The subfamily assignment of each identified LRR-containing protein was deduced from its domain structure. Proteins were classified into the LRR-RLK subfamily if they contained LRRs and a kinase domain, and sometimes other domains such as the malectin, malectin-like, cys-pair and TM domains. Proteins were classified in the NLR subfamily if they included an NB-ARC domain and LRRs, sometimes with a TIR or an RPW8 domain. The LRR-RLP subfamily included proteins with LRRs plus a TM, malectin, malectin-like and/or cys-pair, or LRR-only structures when at least 10 plant-specific LRR repeats were detected. Proteins containing an F-box or an FBD domain in addition to LRRs were classified as F-box-LRR. All other LRR-containing proteins were ranked in the UC group, and for these we performed a BLASTp search with default parameters against the other gene sets (LRR-RLP, LRR-RLK, NLR and F-box) to estimate their probable membership in one of these gene subfamilies. F-box proteins were removed from our datasets and not further considered in the analyses. We ended up with four gene sets: LRR-RLP, LRR-RLK, NLR and UC.

At the end of the construction phase, the complete LRRprofiler pipeline was tested on the manually reviewed *Arabidopsis thaliana* protein dataset downloaded from the Swiss-Prot section (https://www.uniprot.org/) (**Supplemental Methods, Supplemental Figure 5, Supplemental Table 3, Supplemental Data Set 4**) (Boutet et al., 2007) of the UniProt databank (UniProt, 2019). This set was composed of 15,818 sequences. Domain and repeat information was also extracted from the database, especially the number of LRR motifs per sequence and the gene subfamily to which it belonged (LRR-RLP, LRR-RLK, NLR, etc.).

### Rice transcription factor dataset

Transcription factor genes (TF) were identified in the proteome predicted from the three publicly available annotations of the Nipponbare rice reference genome using iTAK (Zheng et al., 2016). Nine subfamilies were considered: C2H2, FAR1, MYB-related, WRKY, NAC, AP2/ERF-ERF, bHLH, bZIP and MYB.

### Annotation transfer from Nipponbare to Kitaake

The first phase consisted of localization of regions of interest in the Kitaake genome, i.e. region homologous to Nipponbare LRR-CR loci. Nipponbare LRR-CR protein sequences from our expert annotations were aligned to the Kitaake genome with tBLASTx (Altschul et al., 1990). Only high scoring pairs (HSPs) with more than 65% identity with Nipponbare LRR-CR protein fragments and spanning at least 50% of the Nipponbare query protein were retained. To define coherent candidate regions in Kitaake, HSPs from the same Nipponbare query protein had to be located less than 5,000 bp apart, except when the query Nipponbare homologous gene had a longer intron. In that case, the Nipponbare intron length plus 500 bp was used as the upper bound for the distance separating Kitaake HSPs. Multiple regions of interest could be found for a single Nipponbare protein. This allowed us to annotate genes duplicated in the Kitaake genome even if a single gene copy was present in the Nipponbare genome. In a second phase, gene model determination was attempted in three consecutive steps for each region of interest. Only regions that could not be successfully annotated at a given step passed to the next one. In the first step, the Nipponbare query exons are mapped to the target Kitaake region of interest with BLASTn. A gene model was then reconstructed based on ordered HSPs. The gene model reconstruction quality was checked by comparing the predicted protein with that of Nipponbare using BLASTp. The gene model was retained if all expected exons were present and the Kitaake protein sequence had more than 90% identity with the Nipponbare protein sequence. Otherwise annotation of this region was delegated to the second step. In the second step, the Exonerate cdna2genome model (Slater and Birney, 2005) was run independently for every remaining query/target pair. The Exonerate output GFF file was parsed to construct the target gene model and to document putative frameshift positions. Again, the Kitaake predicted protein was compared to the Nipponbare query sequence with BLASTp and retained if the coverage and identity were above 90% and 75%, respectively. Otherwise annotation of this target region was delegated to the third step. In the third step, the remaining loci were reconstructed with the Exonerate protein2genome model. This model is better at finding the correct reading frames when the target and model loci are more divergent, but it fails to correctly annotate type 1 and 2 splicing sites (intron/exon junction falling inside a codon). This problem arises since it uses the same reading frame to translate the whole genomic sequence (the six reading frames are tested, but each resulting translation just uses one of them). To overcome this issue, intron junctions are then corrected with a Python script that looks for canonical splicing sites in a range of two nucleotides before and after the current junctions. Finally, gene models highly divergent from the Nipponbare query sequence, with multiple premature stop codons or without start or terminal stop codons, and overlapping frameshifts are tagged to be manually checked.

### Identification of alleles between Nipponbare and Kitaake

We used Synmap (Lyons and Freeling, 2008) to identify LRR-CR allelic pairs, i.e. genes having the exact same chromosomic position in Nipponbare and Kitaake. Synmap was developed to identify orthologous genes between different species based on microcolinearity conservation, and it identifies blocks of genes of conserved order and position. It retrieves a list of relationships between genic repertoires of two genomes. We identified alleles by first selecting genes for which Synmap found a reciprocal relationship, i.e. a relationship found in both Nipponbare to Kitaake and Kitaake to Nipponbare comparisons. Genes for which allelic relationships could not be unambiguously resolved by Synmap were manually resolved, when possible, using VISTA (Mayor et al., 2000) and Artemis (Carver et al., 2012) software packages.

### Availability of new reannotated data

A new identifier was allocated to each LRR-CR gene unravelled by this procedure. These identifiers use the <OSJnip_ChrXX_00000000> or <OSJkit_ChrXX_00000000> pattern for Nipponbare and Kitaake loci, respectively, with XX being the chromosome number followed by the start codon position of the coding sequence (CDS) on the chromosome (**Supplemental Data Sets 1 and 2**).

According to the MACSE convention (Ranwez et al., 2011; Ranwez et al., 2018), indels causing frameshift mutations have been pinpointed by the presence of one or two ‘!’ characters in the nucleotide and protein sequences of non-canonical genes and are available in an additional specific dataset posted on the website (https://rice-genome-hub.southgreen.fr/content/geloc).

## Supplemental Data

### Supplemental Methods

Validation of the LRRprofiler pipeline.

**Supplemental Data Set 1**. LRR-CR loci from the Nipponbare rice reference genome.

Manually curated annotation of Nipponbare LRR-CR loci. For each LRR-CR gene, the corresponding identifier (ID) from public annotations (IRGSP, MSU and NCBI) are given. Known gene names (aliases) are also given and were retrieved from the NBRP OryzaBase database. Substantial gene information is provided, including the gene sub-family, gene category (canonical or non-canonical), the fact that the gene model was modified (y) or not (n) during the curation process and the number of LRR motifs annotated by the LRR profiler pipeline.

**Supplemental Data Set 2**. LRR-CR loci from the rice KitaakeX genome.

Manually curated annotation of Kitaake LRR-CR loci after transfer of Nipponbare loci. For each LRR-CR gene, the corresponding identifier (ID) from public annotation are given. Substantial gene information is provided, including the gene sub-family, gene category (canonical or non-canonical) and the number of LRR motifs annotated by the LRR profiler pipeline.

**Supplemental Data Set 3**. Allelic relationship and cDNA identity between Nipponbare and Kitaake LRR-CR loci.

Each line provides the allelic pair identifiers (ID) from Nipponbare (NIP) and Kitaake (KIT) cultivars. Allelic relationships were determined with Synmap, followed by manual curation.

**Supplemental Data Set 4**. LRRprofiler results in the Swiss-Prot *A. thaliana* dataset.

LRR-containing proteins extracted from the Swiss-Prot database and results of the LRRprofiler pipeline. The orange section presents information on gene sub-families and number of annotated LRR motifs extracted from the Swiss-Prot database. The blue section presents the LRRprofiler results for each gene.

**Supplemental Figure 1**. Comparison of predicted peptide lengths between Nipponbare publicly available annotations (IRGSP, MSU, NCBI) for LRR-CR and TF loci.

The plots represent pairwise comparaisons of peptide lengths (in amino acids) predicted by IRGSP and MSU (left), IRGSP and NCBI (centre) and MSU and NCBI (right) for LRR-CR genes (upper) and TF genes (lower). Numbers at the bottom right of each graph are the coefficient of determination (R^2^) and gene family average relative length difference (RLD). The grey dotted lines on each graph represent the diagonal y=x (dots are shown along this line when the two considered annotations predicted peptides of the exact same length for a given gene).

**Supplemental Figure 2**. Number of domains and motifs identified with LRRprofiler for the Nipponbare proteomes predicted by publicly available and manualy curated annotations.

**Supplemental Figure 3**. LRR motif number conservation between Nipponbare and Kitaake LRR-CR loci depending on the compared annotation.

The public annotation for the Kitaake genome was compared to the Nipponbare IRGSP (green), MSU (orange) and NCBI (blue) annotations. Our manually curated annotations compared for Nipponbare and Kitaake are shown in pink. The ‘Mean(diff)’ statistic is the average difference in the number of LRR motifs between the two cultivar alleles. The dashed line on each graph represents the y=x diagonal.

**Supplemental Figure 4**. Schematic representation of the LRRprofiler pipeline.

**(A)** HMM profile refinement sub-process for LRR motifs. In a first step, candidate proteins were identified for LRR-RLK or NLR subfamilies respectively using iTAK or hmmsearch with the PF00931 NB-ARC profile. In a first round of LRR identification, one of the two candidate protein sets was run with the SM00370 LRR profile as input. Then identified LRRs were aligned with mafft and a new profile was built with hmmbuild. This new profile was used to search LRR motifs in the chosen candidate protein set. The last steps, i.e. LRR motif alignment, construction of a new profile and searchs for LRR motifs with the new profile, were repeated until the stop condition was fulfilled. Finally, the best profile from all iterations was retrieved as the new specific profile for the subfamily depending on the candidate protein set (NLR or LRR-RLK).

**(B)** LRR-CR protein annotation and classification. Newly refined and SMART LRR profiles (SM00367, SM00368, SM00369, and SM00370) were used to identify LRR-containing proteins from a proteome and to annotate LRR motifs in these proteins, which required handling of motif match overlaps. Other functional domains were annotated with public HMM profiles. For each LRR-containing protein, the gene subfamily (LRR-RLK, LRR-RLP, NLR or F-box-LRR) to which it belonged was deduced from its domain structure. Proteins with at least an NB-ARC and LRRs were NLRs, proteins with at least a kinase and LRRs were LRR-RLKs, while proteins with LRRs and a TM or a malectin(-like) domain without kinase were LRR-RLPs. Structures with an F-box or FBD were F-box-LRRs. All proteins with alternative structures were pooled in the unclassified (UC) group. *F-box and FBD are domains specific to F-box-LRR proteins. F-box-LRRs are other intracellular receptors that were not studied here. Their identification allowed us to separate them from the other LRR-CR proteins.

**Supplemental Figure 5**. Comparison of expected and predicted LRR motifs in protein sequences from the Swiss-Prot *A. thaliana* dataset using publicly available and refined HMM profiles.

Number of expected (x axis) and predicted (y axis) LRR motifs in protein sequences from the Swiss-Prot *A. thaliana* dataset using public and new refined HMM profiles. The upper section compares motif numbers for LRR-RLP and LRR-RLK (concatenated in LRR-RL), and the lower section compares motif numbers for NLR.

**Supplemental Table 1**. Contingency table of canonical/non-canonical and modified/not modified LRR-CR loci from the Nipponbare manually curated annotation.

**Supplemental Table 2**. Percentage of cDNA identity between Nipponbare and Kitaake alleles according to gene sub-families and categories. Categories are given as Nipponbare/Kitaake. C, canonical; NC, non-canonical.

**Supplemental Table 3**. Performance of publicly available and refined LRR HMM profiles in the Swiss-Prot *A. thaliana* dataset.

The third line (Expected) presents the number of proteins belonging to each sub-family and the number of motifs annotated for these proteins in the Swiss-Prot database.

## Acknowledgments

This work was partly supported by the CGIAR Research Program on Rice (CRP RICE) (to AD) and by a PhD fellowship from *Institut Agro* and CIRAD (to CG). Computational resources were provided by the South Green bioinformatics platform. The authors would like to thank Dr. A. Cenci for critical reading of the manuscript.

## Author contributions

CG, NC, AD, CP and VR designed the research; CG, NC, AD performed the research; CG, NC, AD, VR, MS, GD contributed to new analytic and computational tools; CG, NC, AD analyzed the data; CG, NC, AD, VR wrote the paper.

