## Supplemental_Figures for "A New ‘Comprehensive’ Annotation of Leucine-Rich Repeat-Containing Receptors in Rice"

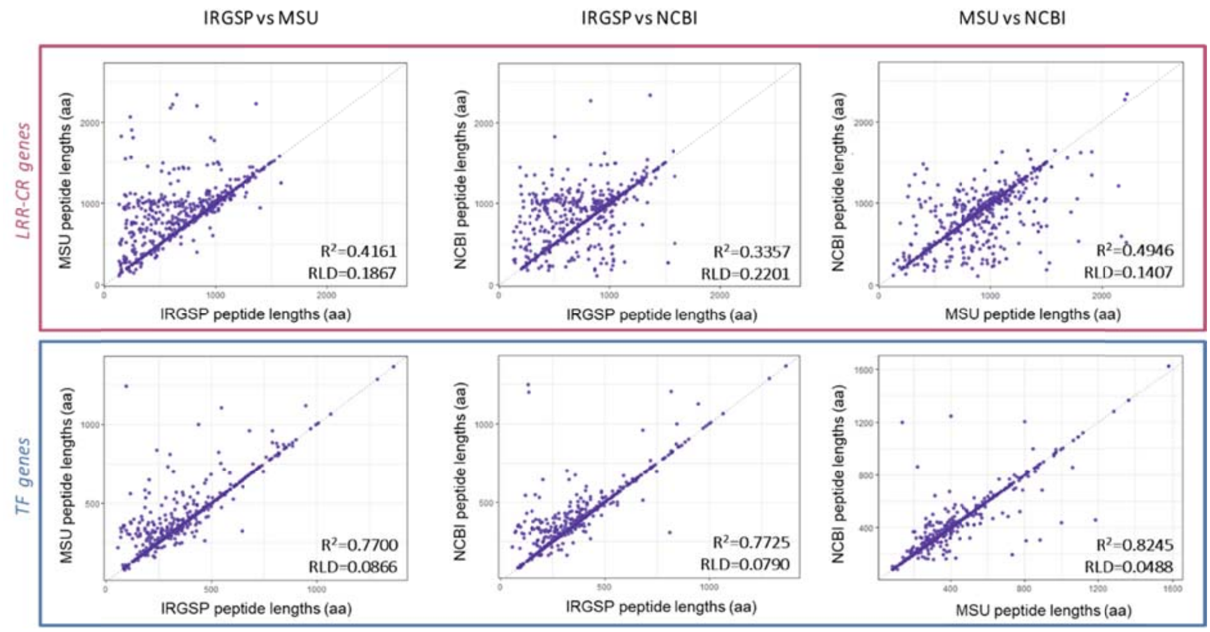


**Supplemental Figure 1.** Comparison of predicted peptide lengths between Nipponbare publicly available annotations (IRGSP, MSU, NCBI) for LRR-CR and TF loci.

The plots represent pairwise comparaisons of peptide lengths (in amino acids) predicted by IRGSP and MSU (left), IRGSP and NCBI (centre) and MSU and NCBI (right) for LRR-CR genes (upper) and TF genes (lower). Numbers at the bottom right of each graph are the coefficient of determination ($R^2$) and gene family average relative length difference (RLD). The grey dotted lines on each graph represent the diagonal y=x (dots are shown along this line when the two considered annotations predicted peptides of the exact same length for a given gene).

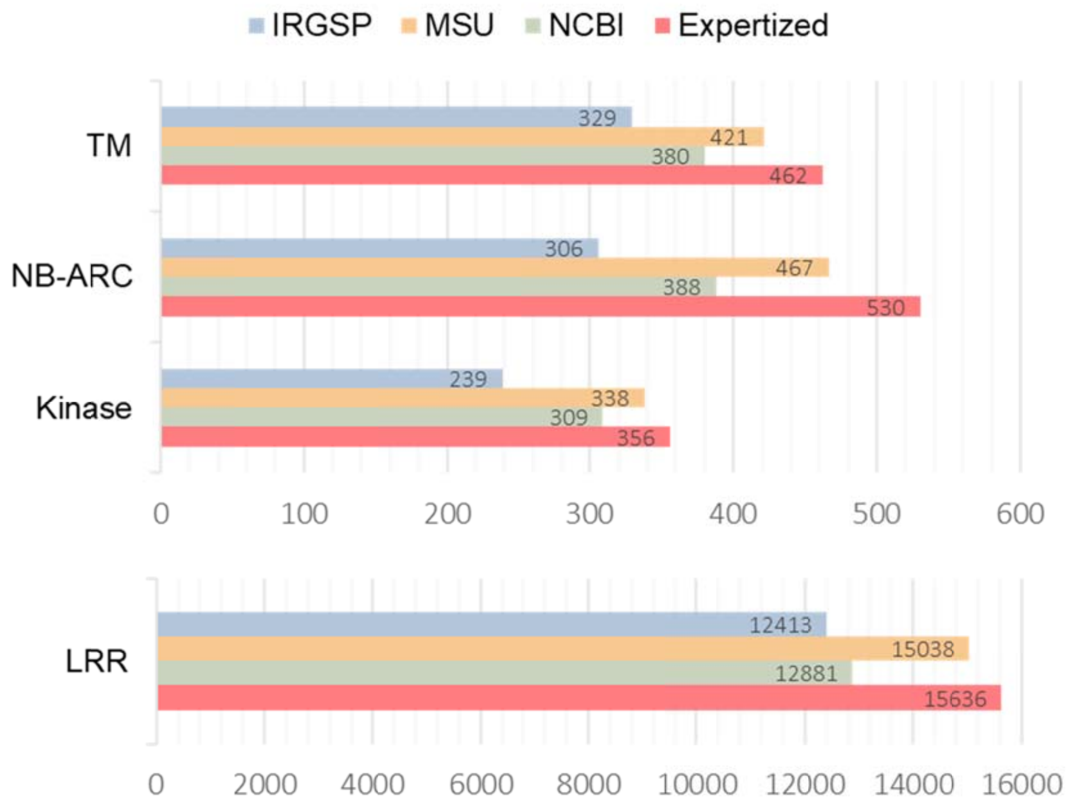


**Supplemental Figure 2.** Number of domains and motifs identified with LRRprofiler for the Nipponbare proteomes predicted by publicly available and manualy curated annotations.

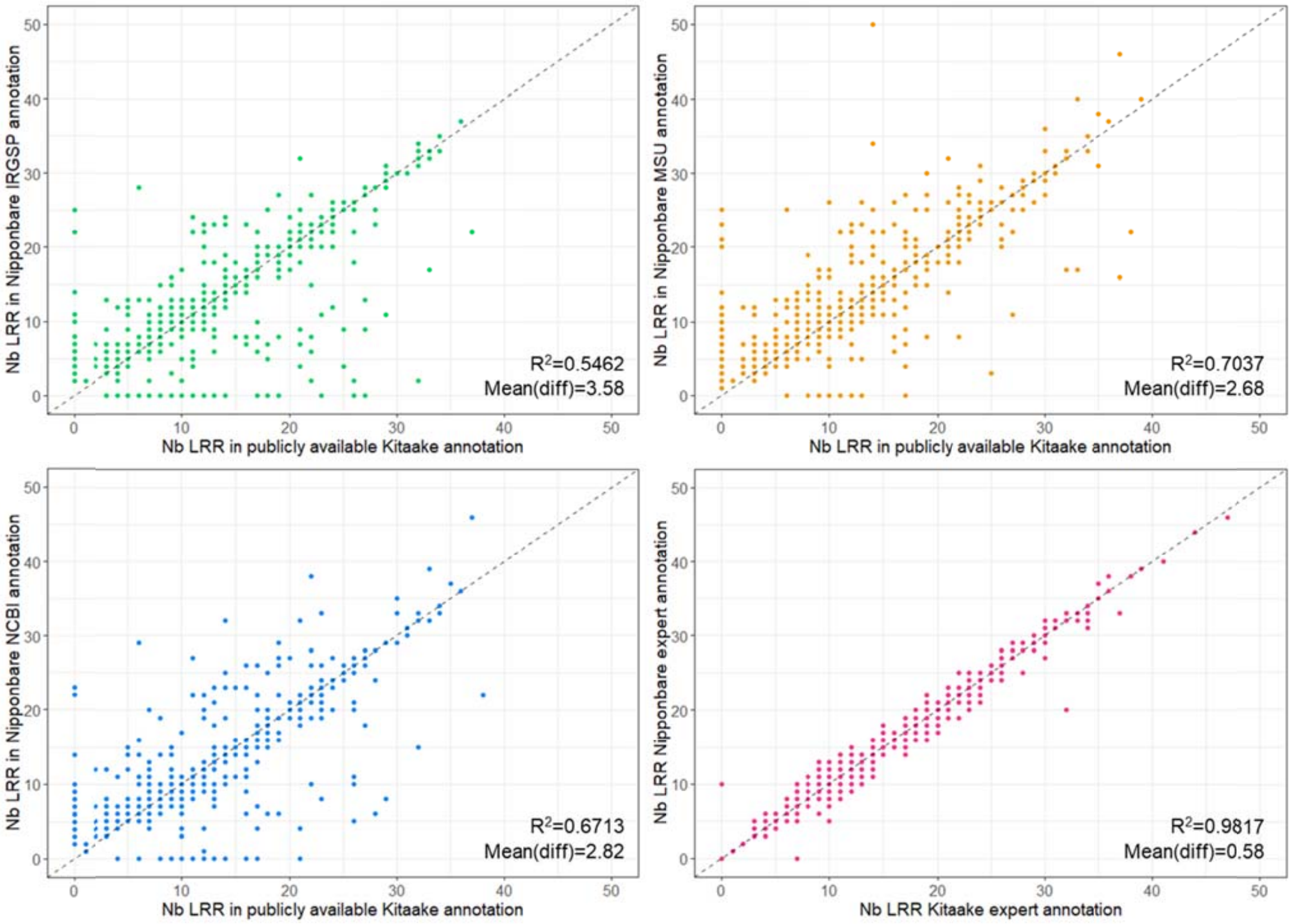


**Supplemental Figure 3.** LRR motif number conservation between Nipponbare and Kitaake LRR-CR loci depending on the compared annotation.

The public annotation for the Kitaake genome was compared to the Nipponbare IRGSP (green), MSU (orange) and NCBI (blue) annotations. Our manually curated annotations compared for Nipponbare and Kitaake are shown in pink. The 'Mean(diff)' statistic is the average difference in the number of LRR motifs between the two cultivar alleles. The dashed line on each graph represents the y=x diagonal.

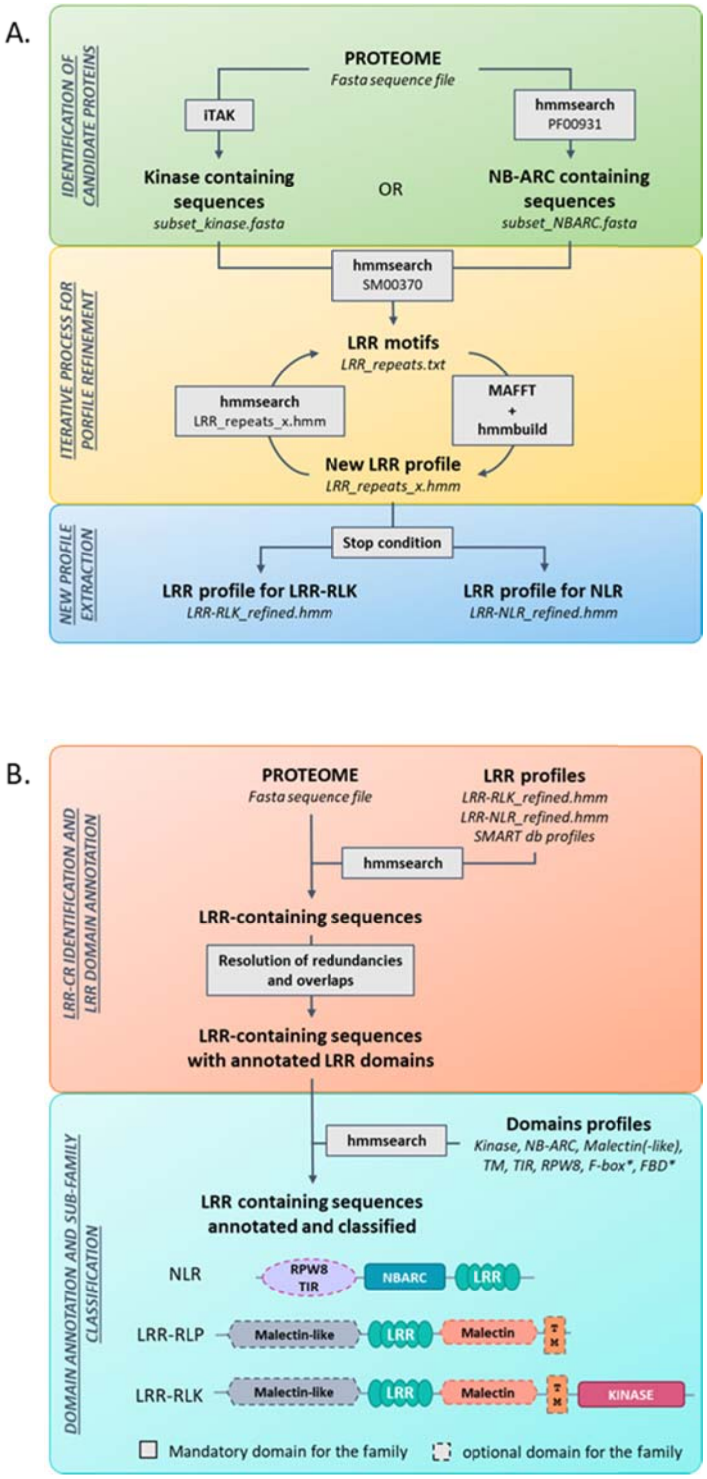


**Supplemental Figure 4**. Schematic representation of the LRRprofiler pipeline.

**(A)** HMM profile refinement sub-process for LRR motifs. In a first step, candidate proteins were identified for LRR-RLK or NLR subfamilies respectively using iTAK or hmmsearch with the PF00931 NB-ARC profile. In a first round of LRR identification, one of the two candidate protein sets was run with the SM00370 LRR profile as input. Then identified LRRs were aligned with mafft and a new profile was built with hmmbuild. This new profile was used to search LRR motifs in the chosen candidate protein set. The last steps, i.e. LRR motif alignment, construction of a new profile and searchs for LRR motifs with the new profile, were repeated until the stop condition was fulfilled. Finally, the best profile from all iterations was retrieved as the new specific profile for the subfamily depending on the candidate protein set (NLR or LRR-RLK).

**(B)** LRR-CR protein annotation and classification. Newly refined and SMART LRR profiles (SM00367, SM00368, SM00369, and SM00370) were used to identify LRR-containing proteins from a proteome and to annotate LRR motifs in these proteins, which required handling of motif match overlaps. Other functional domains were annotated with public HMM profiles. For each LRR-containing protein, the gene subfamily (LRR-RLK, LRR-RLP, NLR or F-box-LRR) to which it belonged was deduced from its domain structure. Proteins with at least an NB-ARC and LRRs were NLRs, proteins with at least a kinase and LRRs were LRR-RLKs, while proteins with LRRs and a TM or a malectin(-like) domain without kinase were LRR-RLPs. Structures with an F-box or FBD were F-box-LRRs. All proteins with alternative structures were pooled in the unclassified (UC) group. *F-box and FBD are domains specific to F-box-LRR proteins. F-box-LRRs are other intracellular receptors that were not studied here. Their identification allowed us to separate them from the other LRR-CR proteins.

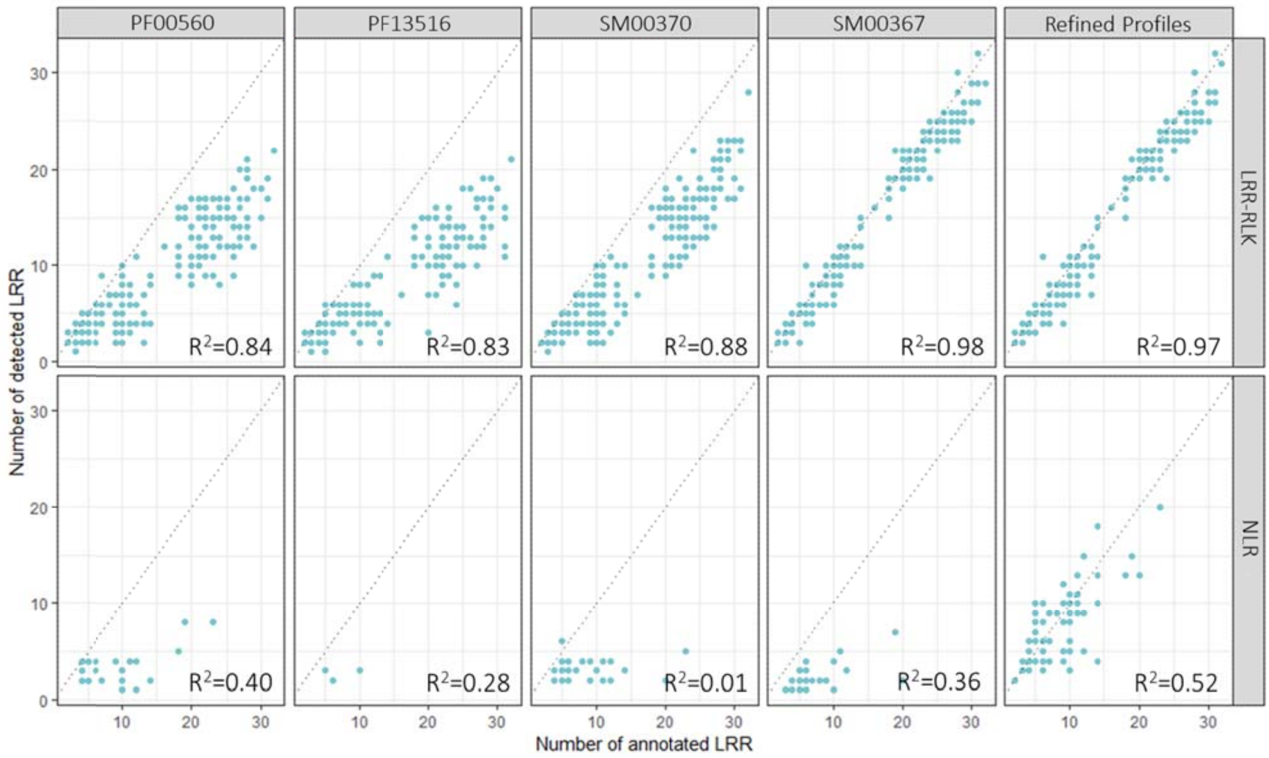


**Supplemental Figure 5.** Comparison of expected and predicted LRR motifs in protein sequences from the Swiss-Prot *A. thaliana* dataset using publicly available and refined HMM profiles.

Number of expected (x axis) and predicted (y axis) LRR motifs in protein sequences from the Swiss-Prot *A. thaliana* dataset using public and new refined HMM profiles. The upper section compares motif numbers for LRR-RLP and LRR-RLK (concatenated in LRR-RL), and the lower section compares motif numbers for NLR.
