## Supplemental_methods for "A New ‘Comprehensive’ Annotation of Leucine-Rich Repeat-Containing Receptors in Rice"

**Supplemental Methods. Validation of the LRRprofiler pipeline**

To automatically detect LRR-CRs in a proteome and identify subfamilies to which they belong (LRR-RLK, LRR-RLP or NLR), we developed the dedicated LRRprofiler pipeline. This pipeline relies on domain detection based on HMM profiles (**Supplemental Figure 4**). It starts by building LRR HMM profiles specific to the considered proteome for two LRR-CR gene subfamilies, i.e. LRR-RLK and NLR (**Supplemental Figure 4A**). Then it uses the two refined LRR profiles, in addition to others, to detect LRR-containing proteins in the considered proteome and to classify LRR-CR proteins into subfamilies based on protein domain arrangements (**Supplemental Figure 4B**). Sequences that cannot be classified in any subfamily based on their domain structure are tagged as UC (unclassified).

To test the different steps of the pipeline, we used the manually reviewed *A. thaliana* protein dataset extracted from the Swiss-Prot database (Boutet et al., 2007). This protein dataset provided us with a highly reliable reference to which automatic LRR annotation results could be compared. This reference set consists of 15,818 proteins. Among these, 318 proteins belong to the three LRR-CR subfamilies with at least one LRR motif (171 LRR-RLKs, 61 LRR-RLPs, and 86 NLRs. **Supplemental Data Set 4**).

1. **Validation of the LRR HMM profiles obtained in the first LRRprofiler steps**

We compared the new refined profiles for LRR motifs to those publicly available on the Pfam and SMART databases. Four LRR profiles from Pfam (PF00560 (LRR_1), PF07723 (LRR_2), PF07725 (LRR_3), and PF13516 (LRR_6)) and four SMART profiles (SM00370 (LRR), SM00369 (LRR_TYP), SM00367 (LRR_CC), and SM00368 (LRR_RI)) were used to search LRRs on the *A. thaliana* dataset mentioned above. Note that in order to precisely detect and localize each LRR motif, only single motif LRR profiles were used, i.e. available LRR profiles based on multiple tandem LRR motifs were not tested. The hmmsearch software package (HMMER, (Eddy, 2011)) was used to identify sequences matching each of these eight HMM profiles. For each tested profile, we retrieved the number and percentage of both detected proteins and LRR motifs in LRR-RLK, LRR-RLP and NLR (**Supplemental Table 3**).

The publicly available profiles tested gave very heterogeneous results. The percentage of LRR-RLK and LRR-RLP protein recovered by the eight public profiles ranged from 0 (PF07725) to 100% (SM00367). Identifying LRR proteins was easier than identifying all of their LRR motifs, e.g. the profile PF00560 (LRR_1) enabled correct identification of 99.4% of the LRR-RLK proteins but only 61.4% of their motifs. This result is illustrated in **Supplemental Figure 5**, which highlights that with most profiles, although LRR-CR proteins were identified, several LRR motifs were missed. Most importantly, although some of these profiles provided a good annotation of LRR-RLK and LRR-RLP proteins and their LRR motifs (with the best results being obtained with the SM00367 profile that recovered 100% of the expected proteins), all of them performed very poorly for NLR (**Supplemental Table 3** and **Supplemental Figure 5**). At best, only 40.7% of the Arabidopsis NLR proteins were identified (SM00367 profile) and no more than 16.5% of their LRR motifs were found (SM00368 profile).

New profiles were refined by using the SM00370 profile as the initial state. The refinement performed with the LRR-RLK candidate set enabled validation of the strategy by comparing the performance of the new LRR-RLK profiles to that of SMART SM00367 (LRR_CC). The results obtained for these two profiles showed that they had very similar performances to the identified and annotated LRR-RLK and LRR-RLP proteins (100% of LRR-RLK and LRR-RLP identified and 97.9% and 92% of their motifs annotated, respectively; **Supplemental Table 3** and **Supplemental Figure 5**). The new profile obtained for NLR LRR motifs greatly improved the protein and motif detection for NLR genes. It detected 98.8% of NLR proteins and 88.2% of their expected motifs, representing a protein and LRR motif detection gain of 58.1% and 71.7%, respectively.

### 2. The entire LRRprofiler pipeline tested on Arabidopsis training datasets

The entire LRRprofiler pipeline was tested on the manually reviewed *A. thaliana* protein dataset cited above. It was able to identify 100% of all expected proteins for the three subfamilies and to correctly classify 100% of the LRR-RLKs, 100% of the NLRs and 95% of the LRR-RLPs (58 out of 61). The three remaining LRR-RLP sequences fell into the UC subgroup. The pipeline precision was also good as all of the sequences labelled as LRR-RLK, LRR-RLP or NLR were well classified.

Moreover, four genes not included initially in the NLR reference set (no LRR motifs annotated in the Swiss-Prot database) were recovered by our pipeline, thus confirming the LRRprofiler pipeline efficiency (**Supplemental Data Set 4**).

A second test was performed by running the LRRprofiler pipeline on the TAIR10 data. The results were compared to the expert data from Meyers et al. (Meyers et al., 2003). The authors retrieved 147 *A. thaliana* NLR genes containing LRR motifs. Only eight of them were not identified by LRRprofiler. Three were absent from the downloaded TAIR10 dataset (AT4G14610; AT3G25515; AT5G40920) and four were present but lacking the LRR domain (AT3G15700; AT5G66630; AT4G09430; AT5G46490) and corresponded to sequences manually modified in Meyers et al. (2003) The last unidentified sequence (AT4G12020) was an uncommon sequence with EGF and WRKY domains associated with an irregular LRR domain.

These validations showed that the LRRprofiler pipeline built suitable refined LRR HMM profiles, and was very efficient in identifying LRR-CR proteins and LRR motifs in a given proteome. The LRRprofiler pipeline is available for download at https://github.com/cgottin/LRRprofiler.

**Boutet, E., Lieberherr, D., Tognolli, M., Schneider, M., and Bairoch, A.** (2007). UniProtKB/Swiss-Prot. Methods Mol Biol **406,** 89-112.

**Eddy, S.R.** (2011). Accelerated Profile HMM Searches. PLoS Comput Biol **7,** e1002195.

**Meyers, B.C., Kozik, A., Griego, A., Kuang, H., and Michelmore, R.W.** (2003). Genome-wide analysis of NBS-LRR-encoding genes in Arabidopsis. Plant Cell **15,** 809-834.
