## Supplemental_Tables for "A New ‘Comprehensive’ Annotation of Leucine-Rich Repeat-Containing Receptors in Rice"

**Supplemental Table 1.** Contingency table of canonical/non-canonical and modified/not modified loci from the Nipponbare manually curated annotation.

| | **Modified** | **Not modified** | **New** | **Total** |
|---|---|---|---|---|
| **Non-canonical** | 273 | 25 | 7 | 305 |
| **Canonical** | 54 | 692 | 1 | 747 |
| **Total** | 327 | 717 | 8 | 1052 |

**Supplemental Table 2**. Percentage of cDNA identity between Nipponbare and Kitaake alleles according to gene sub-families and categories. Categories are given as Nipponbare/Kitaake. C, Canonical; NC, non-canonical.

| **Nipponbare/Kitaake** | **LRR-RLK** | **LRR-RLP** | **NLR** | **UC** | **Total** |
|---|---|---|---|---|---|
| C/C | 99.76 | 99.39 | 99.07 | 99.99 | 99.43 |
| NC/NC | 99.35 | 97.11 | 98.88 | 99.51 | 98.73 |
| C/NC | 98.47 | 97.39 | 95.69 | 99.06 | 96.96 |
| NC/C | 89.52 | 72.49* | 91.58 | NA | 87.37 |
| Total | 99.56 | 97.91 | 98.72 | 99.77 | 98.95 |

* Concerns three pairs of loci including one pair with a truncated loci in Kitaake genome that lead to only 39% of cDNA identity between pair members.

**Supplemental Table 3.** Performance of publicly available and refined LRR HMM profiles on the Swiss-Prot *A. thaliana* dataset. The third line (Expected) presents the number of proteins belonging to each sub-family and the number of motifs annotated for these proteins in the Swiss-Prot database.

| **Gene Subfamily** | | **LRR-RLK** | | | | **LRR-RLP** | | | | **NLR** | | | |
|---|---|---|---|---|---|---|---|---|---|---|---|---|---|
| **Expected (Swiss-Prot)** | | **Proteins** | | **LRR Motifs** | | **Proteins** | | **LRR Motifs** | | **Proteins** | | **LRR Motifs** | |
| | | **171** | **%** | **1941** | **%** | **61** | **%** | **1333** | **%** | **86** | **%** | **689** | **%** |
| **Pfam** | PF00560 (LRR_1) | 170 | 99.4 | 1192 | 61.4 | 61 | 100 | 797 | 59.8 | 29 | 33.7 | 94 | 13.6 |
| | PF07723 (LRR_2) | 21 | 12.3 | 48 | 2.5 | 8 | 13.1 | 21 | 1.6 | 0 | 0.0 | 0 | 0.0 |
| | PF07725 (LRR_3) | 0 | 0 | 0 | 0 | 0 | 0 | 0 | 0 | 32 | 37.2 | 66 | 9.6 |
| | PF13516 (LRR_6) | 129 | 75.4 | 786 | 40.5 | 55 | 90.2 | 665 | 49.9 | 6 | 7.0 | 14 | 2.1 |
| **SMART** | SM00370 (LRR) | 164 | 95.9 | 1239 | 63.8 | 61 | 100 | 928 | 69.6 | 28 | 32.6 | 89 | 12.9 |
| | SM00369 (LRR_TYP) | 67 | 39.2 | 279 | 14.4 | 24 | 39.3 | 257 | 19.3 | 1 | 1.2 | 4 | 0.6 |
| | SM00367 (LRR_CC) | 171 | 100 | 1892 | 97.5 | 61 | 100 | 1231 | 92.3 | 35 | 40.7 | 78 | 11.3 |
| | SM00368 (LRR_RI) | 5 | 2.9 | 19 | 1.0 | 6 | 9.8 | 36 | 2.7 | 23 | 26.8 | 114 | 16.5 |
| **Refined Profile** | LRR_RLK | 171 | 100 | 1901 | 97.9 | 61 | 100 | 1227 | 92.0 | 24 | 27.9 | 50 | 7.3 |
| | LRR_NLR | 37 | 21.6 | 116 | 6.0 | 17 | 27.9 | 155 | 11.6 | 85 | 98.8 | 608 | 88.2 |
